# Lysosomal calcium signalling contributes to the acute α-adrenergic response via calcium-stimulated adenylyl cyclase 1 and 8

**DOI:** 10.1101/2024.11.25.625232

**Authors:** Emily Akerman, Rebecca A Capel, Matthew Read, Qianqian Song, Samuel J Bose, Serena Calamaio, Daniel Aston, Andreas Koschinski, Scott Lloyd, Laura Bell, Victoria S Rashbrook, Marco Keller, Franz Bracher, Barry VL Potter, Ilaria Rivolta, Antony Galione, Duncan B Sparrow, Alana Conti, Derek A Terrar, Manuela Zaccolo, Rebecca AB Burton

## Abstract

Inositol trisphosphate (IP_3_), a calcium (Ca^2+^)-mobilizing second messenger, releases Ca^2+^ from the sarcoplasmic reticulum (SR) via IP_3_ receptors and modulates adenylyl cyclase (AC) activity in atrial myocytes. Lysosomes participate in Ca^2+^ homeostasis by mobilising Ca^2+^ in response to Nicotinic Acid Adenine Dinucleotide Phosphate (NAADP). We postulate that both downstream activation of Ca^2+^ sensitive AC (AC1 and AC8) and lysosomal Ca^2+^ signalling in response to IP_3_R activation contribute to atrial myocyte function and pacemaking. Ectopic application of phenylephrine (PE) increased chronotropy and inotropy and this response was reduced in the presence of NAADP pathway inhibitors (BZ-194 and SAN4825) and Bafilomycin A1. PE increased cyclic adenosine 3’-5’ monophosphate (cAMP) activity in neonatal rat atrial myocytes (NRAMs) and this was inhibited by NAADP pathway inhibitors. This inhibition was not observed in neonatal rat ventricular myocytes (NRVMs), revealing specificity of this response to NRAMs. We investigated expression of AC1 and AC8 as a possible explanation to these observations. Genetic perturbation of AC1 and AC8 by double-knockout of *Adcy1* and *Adcy8* in a mouse model showed a decrease in positive chronotropic and inotropic response upon cumulative dose of PE in atrial tissue, reduced PE stimulated amplitude of Ca^2+^ transient in isolated atrial myocytes and presented decreased cytosolic cAMP levels in response to PE in neonatal atrial myocytes that was not inhibited by NAADP pathway inhibitors. Our data identifies a link between NAADP and α-adrenergic signalling pathways in atrial myocytes, highlighting that lysosomal Ca^2+^ is an important component of α-adrenergic stimulation in the cardiac atria and warrants further investigation.

## Introduction

Intracellular calcium (Ca^2+^) release from intracellular organelles is a driving force for normal cardiac pacemaker cell automaticity (Maltsev, Vinogradova et al. 2006). In 1983, Ca^2+^ release induced by inositol 1,4,5-trisphosphate (IP_3_) from the endoplasmic reticulum (ER) was demonstrated in pancreatic acinar cells for the first time (Streb, Irvine et al. 1983). Since then, this mechanism has been shown in many other cell types (Berridge 2009); as well as in non-excitable cells such as proliferating lymphocytes (Feske 2007) and excitable cells including cardiac myocytes (Kockskämper, Zima et al. 2008). IP_3_ signalling enhances Ca^2+^-induced force oscillations (Zhu and Nosek 1991) but compared to the role played by type 2 ryanodine receptors (RyR2) in Ca^2+^ induced Ca^2+^ release (CICR), the involvement of IP_3_ signalling in cardiac cellular Ca^2+^ handling is less well studied. Abnormal cardiac intracellular Ca^2+^ dynamics has been identified as a significant contributor to both ventricular and atrial arrhythmias (Deo, Weinberg et al. 2017). Ca^2+^ signalling is normally highly coordinated in atrial myocytes and abnormal intracellular Ca^2+^ homeostasis is observed in chronic atrial fibrillation (AF) (Nattel and Dobrev 2012). RyR2 are Ca^2+^ release channels found on the sarcoplasmic reticulum (SR) in the heart (Wang, Shi et al.), and have received significant attention over the years. More recently, IP_3_ receptors (IP_3_R) have been identified as a second pathway for agonist-induced Ca^2+^ release and have been gaining significant interest due to the importance of IP_3_-inducing agonists such as angiotensin II, endothelin-1 (ET-1), and norepinephrine (Hund, Ziman et al. 2008).

IP_3_ as a Ca^2+^- mobilizing second messenger has recently been shown to modulate adenylyl cyclase (AC) activity in atrial (Capel, Bose et al. 2021) and sino-atrial node myocytes, with IP_3_R activation leading to increased cyclic adenosine 3’-5’ monophosphate (cAMP) (Bose, Read et al. 2022). IP_3_ has been shown to be positively inotropic in atrial (Lipp, Laine et al. 2000) and ventricular (Nosek, Williams et al. 1986) preparations, and in the sinoatrial node it has been found to be positively chronotropic (Ju, Liu et al. 2011).

Another Ca^2+^ mobilising second messenger is nicotinic acid adenine dinucleotide phosphate (NAADP). NAADP targets lysosome-like acidic compartments (Calcraft, Ruas et al. 2009). Lysosomes have been shown to associate with IP_3_R clusters and sequester Ca^2+^ released by IP_3_R (Atakpa, Thillaiappan et al. 2018), suggesting the involvement of IP_3_Rs in the regulation of Ca^2+^ exchange between the endoplasmic reticulum and lysosomes. Crosstalk between NAADP signalling and that mediated by IP_3_, Ca^2+^, and cyclic ADP ribose (cADPR) exists in many cell types (Churchill and Galione 2001, Boittin, Galione et al. 2002, Yuan, Arige et al. 2024).

Lysosomes express various channels that allow Ca^2+^ release, including Two-Pore Channel 2 (TPC2), Transient Receptor Potential Mucolipin (TRPML), and ATP-regulated P2X4 receptors (Morgan, Platt et al. 2011). Micro-domains allowing for highly localized communication between the SR and lysosomes have been shown to exist in both ventricular and atrial cardiomyocytes (Capel, Bolton et al. 2015, Aston, Capel et al. 2017). The cytosolic Ca^2+^ signals that are evoked by TRPML or TPC2 can be amplified by Ca^2+^ induced Ca^2+^ release (CICR) through IP_3_R or RyR2 when lysosomes are closely opposed with endoplasmic reticulum (Patel, Marchant et al. 2010, Galione 2015) and Ca^2+^ released by SR/ER channels can be rapidly sequestered by lysosomes (Capel, Bolton et al. 2015, Atakpa, Thillaiappan et al. 2018). Lysosomes can interact with other Ca^2+^ storage organelles to increase Ca^2+^ release and influence localized Ca^2+^ signalling microdomains (Morgan, Davis et al. 2013, Aston, Capel et al. 2017).

IP_3_-mediated Ca^2+^ release from the SR can stimulate cytosolic adenosine monophosphate (cAMP) synthesis via Ca^2+^-sensitive adenylyl cyclases AC1 and/or AC8 and is essential for modulation of atrial Ca^2+^ signalling in response to alpha-adrenergic stimulation under physiological conditions (Capel, Bose et al. 2021).

We postulate that IP_3_ and NAADP signalling pathways are involved in the modulation of atrial Ca^2+^ signalling in response to α-adrenergic stimulation under physiological conditions, resulting in the modulation of cAMP synthesis via AC1 and/or AC8. Knowledge relating to the interaction between IP_3_, ACs and NAADP pathways may provide insights concerning mechanisms of atrial arrhythmias and present targets for novel therapies.

## Results

### [1] Lysosomal Ca^2+^ signalling contributes to the acute α-adrenergic response

Spontaneously beating right atrial preparations with intact sinoatrial node (SAN) were used to investigate responses by monitoring changes in spontaneous beating rate during cumulative doses of the α1 agonist, phenylephrine (PE). Figure 1A presents dose response curves for spontaneous beating rate that were generated in response to 0.1 up to 30 μM PE (Landzberg, Parker et al. 1991). Under control conditions, in the presence of vehicle (0.001 % DMSO), the spontaneous beating rate of the right atria increased by a maximum of 23.69 % (19-30.13) at 30 μM PE, with an EC_50_ of 1.4 μM (0.3-5.8) (*n*=7). NAADP inhibitor BZ-194 (Dammermann, Zhang et al. 2009, Zhang, Watt et al. 2018), and Bafilomycin A1, a disruptor of lysosomal function by inhibiting the vacuolar-type H^+^-ATPase pump, reduced the response of beating rate to PE compared to the control condition at all concentrations tested and caused a significant rightward shift in the EC_50_. The presence of 500 μM of BZ-194 significantly reduced the response to 1 µM PE (*P*=0.037), with a maximum increase of 11.01 % (8.6-15.6) at 30 μM (*P*<0.0001) of PE, corresponding to a reduction of 53.5 % compared to control, with an EC_50_ of 3 µM (0.7-14.4) (n=4, *P*<0.001). Incubation with 100 nM Bafilomycin A1 caused a maximum increase of 9.42 % (6.4-22.7) at 30 μM PE (*P*<0.0001), a significant reduction compared to control (60.2 %), with a rightward shift of EC_50_ to 3.1 µM (0.4-50.5) (*n*=5, *P*<0.0002). Similarly, the presence of 10μM of SAN4825, an ADP-ribosyl cyclase inhibitor that reduces NAADP synthesis, also significantly reduced the beating rate response to PE from 1 µM (*P*<0.0001), with a maximum increase of 7 % (4.8-12) at 30 μM PE (*P*<0.0001), corresponding to a reduction of 70.4 % compared to control, with an EC_50_ of 2.9 µM (0.4-24.4) (*n*=6, *P*<0.0001). No significant differences were seen in beating rate before after addition of inhibitory drugs or before addition of PE (Figure 1B and Supplementary Figure 1A).

**Figure 1:**
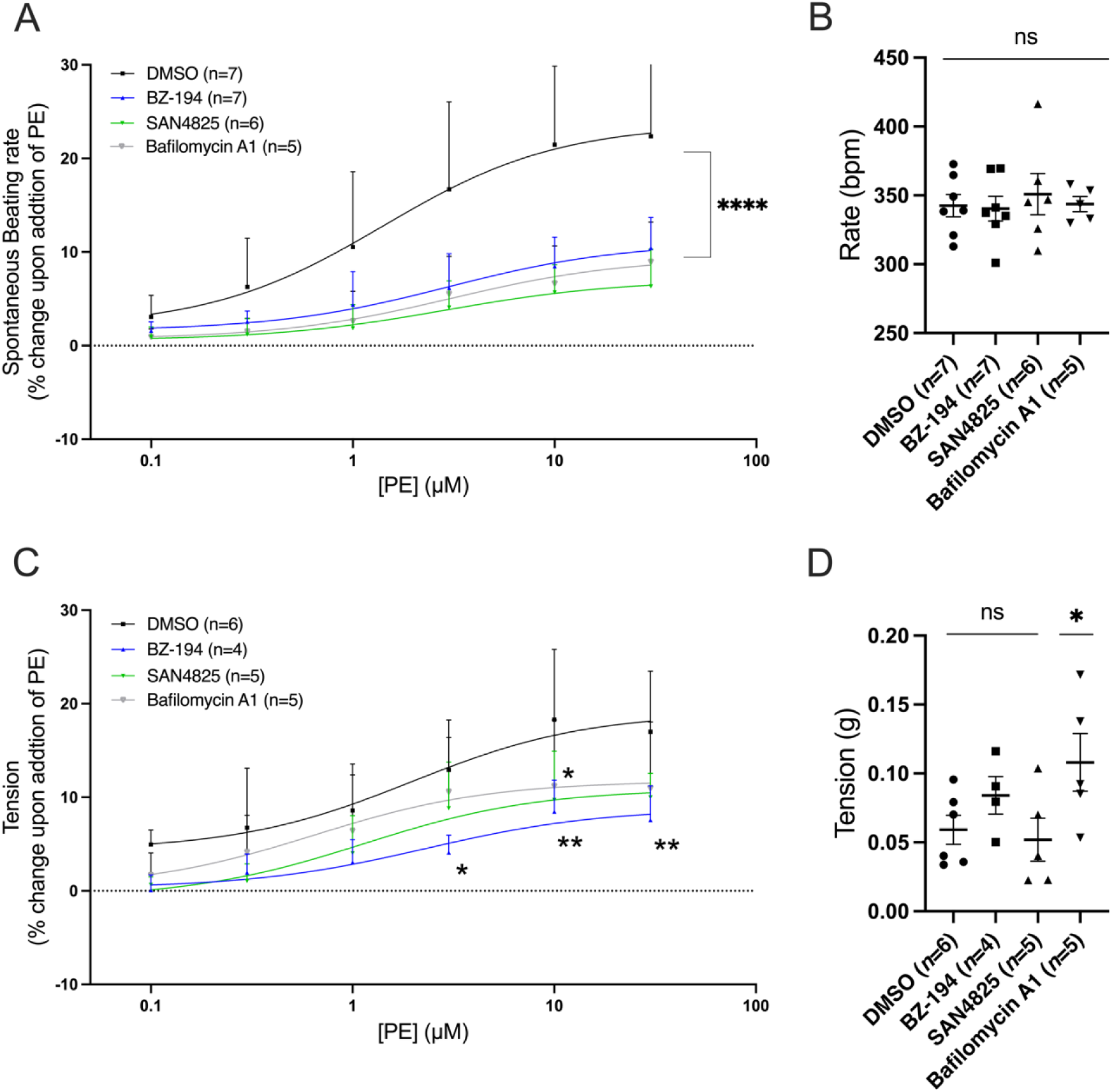
Inhibition of NAADP pathway reduces the positive chronotropic and inotropic effect of α-adrenergic stimulation. **A:** Rate responses to PE (0.1 µM-30 µM) in spontaneously beating murine atrial preparations under control conditions (*n*=7, black) and in the presence of 500 μM BZ-194 (*n*=7, blue), 10 μM SAN4825 (*n*=6, green) and nM of Bafilomycin A1 (n=4, grey). **B**: Basal beating rate in the presence of 1μM Metoprolol before addition of PE. **C**: Tension (g) responses to PE (0.1 µM-30 µM) in paced (5 Hz) left murine atrial preparations under control conditions (n=5) and in the presence of 500 μM BZ-194 (n=3, blue), 10 μM SAN4825 (n=5, green) and 100 nM of Bafilomycin A1 (n=4, grey). **D**: Basal tension after 1 μM Metoprolol. Dose-response curves were fitted using log(agonist) vs. response (three-parameter model). Asterisks indicate significance level for effect of BZ-194, SAN4825 and Bafilomycin A1 compared to control at individual concentrations, determined using 2-way repeated measures ANOVA followed by Dunnett’s multiple comparisons test. Normality was assessed using the Shapiro-Wilk test. Data on graph are represented as mean ± SEM; *p<0.05; **p<0.01; ****p<0.0001.

Intact left atria were used to record changes in contractile force in response to PE when in the presence of NAADP inhibitors and Bafilomycin A1 with beating rate and voltage held constant (Figure 1C). Under control conditions, the maximum change in tension generated by left atria was 19.1 (14.9-25.5) % at 30 μM PE, with an EC_50_ of 2 (0.3-10.5) μM (*n*=6). Bafilomycin A1 reduced the maximum tension generated by PE at all concentrations tested. The presence of 500 μM BZ-194 significantly reduced the response to the highest concentrations of PE with a maximum increase of 8.8 (6.6-13.3) % at 30 μM of PE (*P*<0.01) with a non-significant change of EC_50_ (2.3 (0.2-14) μM (*n*=4)). Incubation of SAN4825 or Bafilomycin A1 showed a trend to reduction in responses compared to control and only showed a significant decrease at 10 µM PE (*P*<0.05). 10 μM of SAN4825, led to a maximum increase of 10.9 (8.1-14.6) % at 30 μM PE with an EC_50_ of (1.1 (0.2-5.5) μM (*n*=5)). Incubation with 100 nM Bafilomycin A1 caused a maximum increase of 11.8 (8.3-16.9) % at 30 μM PE, with an EC_50_ of 0.6 (0-9.1) μM (*n*=5).

Baseline tension was measure after addition of inhibitory drugs or before addition of PE no significant differences were seen in BZ194 and SAN4825 conditions, however Bafilomycin A1 was higher compared to control (*P<0.05, Figure 1D and Supplementary Figure 1C). The β-adrenoceptor antagonist Metoprolol (1µM) (Capel, Bose et al. 2021) was added to each condition in left and right atria to ensure no confounding effects of β-adrenergic response during PE cumulative doses, no differences in percentage change was observed in beating rate or tension (Supplementary Figure 1B and 1D).

### [2] AC1 and AC8 are implicated in chronotropic and inotropic effect of PE in intact atria

A Double Knock-Out (DKO) transgenic mouse line that lacks expression of both *Acyl1* (AC1) and *Acyl8* (AC8) was kindly supplied from the Conti lab (Moulder, Jiang et al. 2008) and used to study the role of these Ca^2+^-sensitive enzymes in α-adrenergic responses. These animals were previously used to study AC1 and AC8 expression in the brain and their role in hippocampus dependent long-term memory (Wong, Athos et al. 1999). The phenotype of these mice has not previously been investigated outside the brain. We conducted detailed structural analysis to assess whether the knockout of AC1 and AC8 impacted the anatomy of the heart. We conducted 3D imaging of wildtype (WT) (Weninger and Mohun 2007) and DKO hearts using high-resolution episcopic microscopy (HREM). Results showed no significant differences in right (Figure 2Ai) or left atrial (Figure 2Aii) or ventricular (Figure 2Aiii), size. Representative images of a WT heart and a DKO heart can be seen in Figure 2B and 2C and online Supplementary Videos 1 (WT heart) and 2 (DKO heart). Additionally, to no phenotypic changes, there was no difference in the baseline beating rate between WT and DKO in spontaneously beating *ex vivo* intact right atrial preparations (ns; 335 ± 12 vs 354 ± 8 beats.minute^-1^ respectively, P>0.05, n=14, Figure 3B). Beating rate at the start of the protocol and prior to the addition of PE or Metoprolol, in WT and DKO mice were compared, and cause a decrease in all conditions, starting rate of WT group was higher but with no significant differences (Supplement Figure 2.A) but both groups showed no difference in the baseline beating rate after addition of Metoprolol (Figure 3B and Supplement Figure 2B and 2C) causing the WT group to have a greater decrease response to Metoprolol (32 % decrease, n=8) compared to DKO (23.4 %, n=14, *P<0.05, Supplement Figure 2.D).

**Figure 2:**
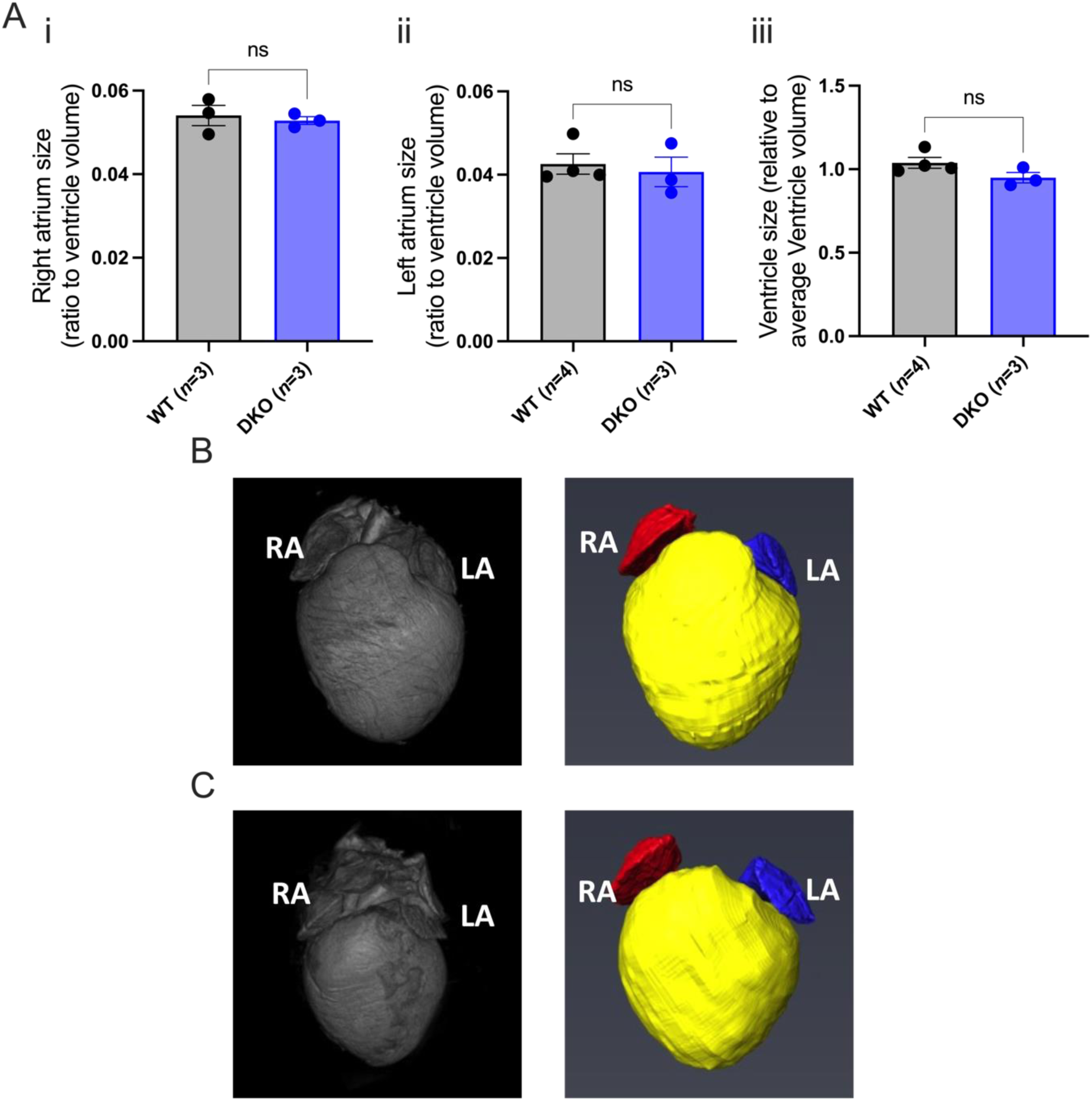
Knockout of AC1 and AC8 expression shows no phenotypic or morphological changes in the heart. **Ai**: Comparison of right atrium size relative to average ventricle volume in WT (n=3) and DKO (n=3) mice hearts. **ii**: Comparison of left atrium size relative to average ventricle volume in WT(n=4) and DKO (n=3) mice hearts. **iii**: Comparison of ventricle size relative to average ventricle volume in WT (n=4) and DKO (n=3) mice hearts. **B**: Representative image of WT heart. **C**: Representative image of DKO heart. Data are represented as mean ± SEM and were analysed using paired t-test; non-significant (ns)=P>0.05.

**Figure 3:**
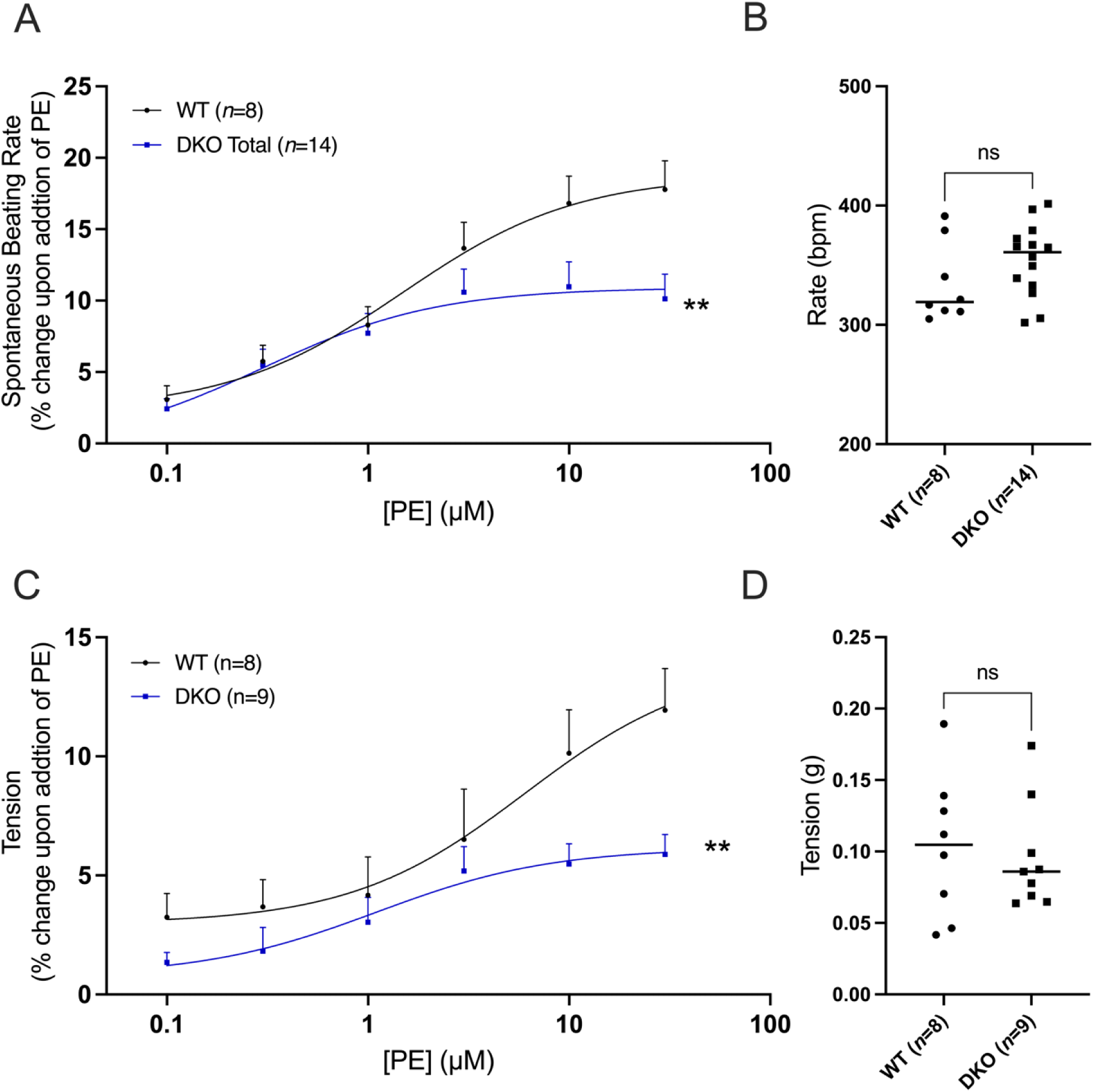
AC1 and AC8 expression impacts positive chronotropic and inotropic responses to α-adrenergic stimulation in atria. **A**: Dose response curves to show the change in beating rate on cumulative addition of PE (0.1 μM-30 μM) to spontaneously beating mouse right atria preparations in WT mice (black, n=8) and DKO mice (blue, n=14). **B**: Comparison of spontaneously baseline beating rate of WT(n=8) and DKO (n=14) C57s murine right atrial preparations in PSS in the presence of 1 µM Metoprolol. **C**: Dose response curves to show the force of contraction on cumulative addition of PE (0.1μM-30μM) to spontaneously beating mouse right atria preparations in WT mice (black, n=8) and DKO mice (blue, n=9). **D**: Comparison of baseline tension of WT (n=8) and DKO (n=10) C57s murine right atrial preparations in PSS after addition of 1 µM Metoprolol. Dose-response curves were fitted using log(agonist) vs. response (three-parameter model). Asterisks indicate significance level for effect of DKO compared to WT at individual concentrations as determined using 2-way repeated measures ANOVA followed by Šídák’s multiple comparisons test; **P<0.01. Data in B and D are represented as mean and were analysed using paired t-test; non-significant (ns)=P>0.05.

Previous published work has used pharmacological tools such as MDL-12330A to inhibit AC activity (Capel, Bose et al. 2021). Here we use the DKO transgenic mouse line to determine the influence of AC1 and AC8 on chronotropic and inotropic responses to α-adrenergic stimulation. Figure 3A presents chronotropic dose-response curves to PE carried out on spontaneously beating isolated murine right atria in the presence of Metoprolol 1 μM. WT mice showed a positive chronotropic response to PE that fits a standard agonist dose-response curve with an EC_50_ of 1.5 (0.5–4.2) μM and a maximum rate increase of 17.8 ± 2 % (n=8, Figure 3B). In the DKO there was a smaller chronotropic response with a maximum rate increase of 10.1 ± 1.7 % (n=14, **P<0.01, Figure 3B) representing a 43 % reduction compared to WT and an EC_50_ of 0.3 (0–1.8) μM.

Intact left atria from WT and DKO mice were used to investigate the influence of AC1 and AC8 on the inotropic dose-response curve to PE (Figure 3C). Dose response curves in Figure 3D show the change in tension generated in response to cumulative additions of PE in the concentration range of 0.1–30 μM. Left atrial preparations were stimulated to beat at a constant rate (5Hz) in the presence of 1 μM Metoprolol. In WT mice, the positive inotropic response to PE had an EC_50_ of 6 μM (1–106.2 μM) and a maximum tension increase of 11.94 ± 1.7 % (n=8). DKO data show a leftward shift in the EC_50_ of 1.1 μM (0.2–8.1 μM) with a significant decrease in maximum tension of 5.9 ± 0.8 % (n=9; **P<0.01), representing a 50.7 % reduction compared to WT. Starting tension of WT and DKO mice were compared and no significant differences were observed between the two groups (ns; 0.1031 +/- 0.02 g, n=8 and 0.0957 +/- 0.09 g, n=9 respectively, P>0.05). Baseline tension after addition of Metoprolol in WT and DKO mice were compared; and no significant differences were observed between the two groups (Supplement Figure 2.E-G).

In the process of backcrossing animals to generate the DKO genotype, 3 groups of animals heterozygous (Het) and homozygous (Hom) for AC1 and AC8 were also generated: [1] Het for both AC1 and AC8 (HetAC1/8), [2] Het for AC1/Hom for AC8 and [3] Hom for AC1/Het for AC8 mice. Rate studies were also conducted on these animals to study the impact separately of either AC1 or AC8, and also presented no changes in beating rate before addition of PE (Supplement Figure 3A). Supplement Figure 3A-C compares dose-response curves to PE between different genotypes created in the concentration range 0.1–30 μM carried out on spontaneously beating isolated murine right atria in the presence of 1 µM Metoprolol. Hom for AC1/Het for AC8 genotype show an EC_50_ of 3.7 (3.2–9.2) μM and a maximum rate increase of 10.9 ± 2.7 % (n=5; *P<0.05; Supplement Figure 3C), the only genotype with a significant decrease in maximum rate response to PE compared to WT with a reduction of 38.7 % in beating rate compared to WT. The effect of Het AC1/8 on inotropy was also investigated and showed no significant change in response to PE but had a significant decrease in the presence of 1 µM Metoprolol (Supplement Figure 3D).

### [3] Lysosomal signalling contributes to an increase in cellular cAMP upon acute α-adrenergic stimulation

Here, we examined the dynamic interactions between α-adrenergic and NAADP signalling pathways and their influence on cAMP activity. This was achieved using cAMP-sensitive FRET experiments in the presence of NAADP inhibitors and PE stimulation in Neonatal Rat Atrial Myocytes (NRAMs) infected with cytosolic FRET biosensor EPAC^SH187^ (Klarenbeek, Goedhart et al. 2015). Protocol is shown in Figure 4A and the presence of lysosomes in NRAMs was confirmed using confocal imaging with labelling of Lysosome-Associated Membrane Protein-2 (LAMP2), shown in Figure 4B.

**Figure 4:**
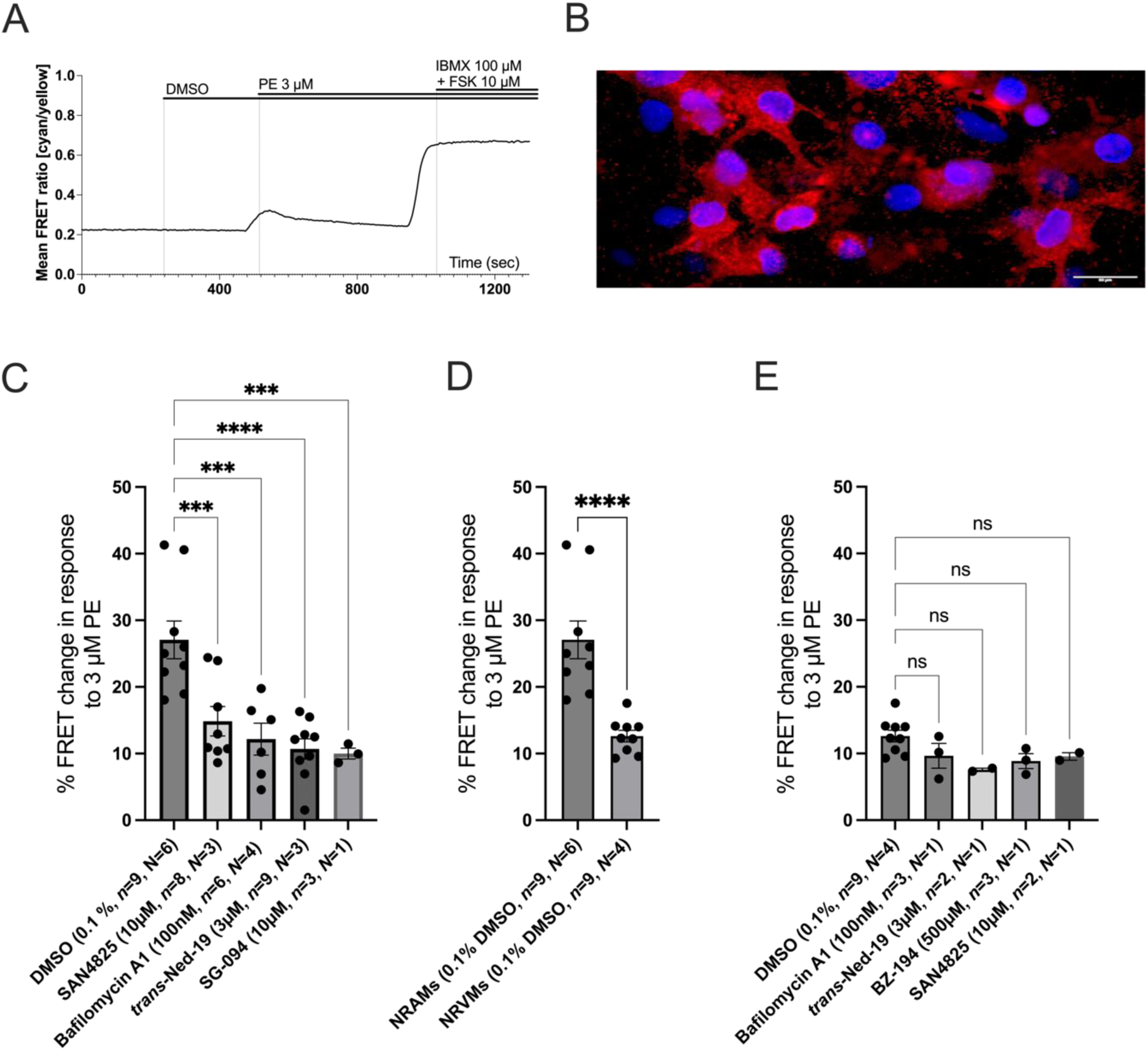
Lysosomal signalling contributes to the increase in cellular cAMP upon acute α-adrenergic stimulation. **A**: FRET Protocol in neonatal myocytes, after 240s of baseline 0.1 % DMSO or drug were added. At 480s 3 µM PE and at 980s saturation response with 10 µM FSK and 100 µM IBMX. **B**: Representative confocal images of fixed cultured NRAMs stained with DAPI (blue) for the nucleus and immunolabelled with LAMP2 (red), scale bar represents 20 µm. **C**: Comparison of peak response to PE, in FRET change (normalised to maximum response) in single NRAMs expressing EPAC-S^H187^ in response to 3 µM PE under control conditions (*n*=9, N=6) and following addition of 10 µM SAN4825 (*n*=8, N=3), 100 nM Bafilomycin A1 (*n*=6, N=4), 10 µM *trans*-Ned-19 (*n*=9, N=3), 10 µM SG-094 (*n*=3, N=1) and 500 µM BZ-194 (*n*=7, N=4). **D**: Comparison of peak response to PE, in FRET change (normalised to maximum response) in single NRAMs (*n*=9, N=6) and in NRVMs (*n*=9, N=6) expressing EPAC-S^H187^ in response to 3 µM PE under control conditions. **E**: Comparison of peak response to PE, in FRET change (normalised to maximum response) in single NRVMs expressing EPAC-S^H187^ in response to 3 µM PE under control conditions (n=9, N=4) and following addition of 100 nM Bafilomycin A1 (n=3, N=1), 1 µM *trans*-Ned-19 (n=2, N=1), 500 µM BZ-194 (n=3, N=1), 10 µM SAN4825 (n=2, N=1). Data are represented as mean ± SEM. Normality was assessed using the Shapiro-Wilk test for panel A and was analysed using 2-way repeated measures ANOVA followed by Dunnett’s multiple comparisons test in panel A, non-parametric Mann-Whitney U test in panel B and panel C was analysed using Kruskal Wallis followed by Dunn’s multiple comparison. n=experiments; N=neonatal isolations. Non-significant (ns), ***P<0.001, ****P<0.0001.

Following a period of stabilisation to establish baseline, 3μM PE was administered and the peak increase in FRET ratio was observed. This peak was expressed as a percentage of a saturating response obtained by the addition of 10 μM FSK and 100μM IBMX. The mean increase in FRET ratio in response to PE was 27.1 ± 2.8 % (n=9, N=6). The addition of inhibitors (10 µM SAN4825, 100 nM Bafilomycin A1, 1 µM trans-Ned-19, 10 µM SG-094 and 500 µM BZ-194) resulted in a significantly attenuated increase in cellular cAMP activity in response to PE under all conditions compared to control (Mean changes shown in Figure 4C and representative FRET traces showing the effect of each individual inhibitor are shown in Supplementary Figure 4A).

These FRET experiments were repeated in Neonatal Rat Ventricular Myocytes (NRVMs) to investigate the specificity of cAMP response to NAADP pathway in response to PE stimulation. Addition of 3μM PE to NRVMs under control conditions in the absence of inhibitor resulted in a FRET change of 12.6 ± 0.8 % (n=9, N=4, Figure 4D). This was significantly lower than the control response seen in NRAMs (P<0.001, Figure 4C). In contrast to NRAMs, the addition of inhibitors to NRVMs resulted in a small inhibition or no significant changes response to PE compared to control (Figure 4E and Supplemental Figure 4B).

### [4] Acute α-adrenergic stimulation leads to lysosomal Ca^2+^ signalling via AC1 and AC8 Activation

Differences in cAMP levels in the presence of NAADP pathway inhibitors in response to PE were next compared in WT and DKO mice (Figure 5). Figure 5A shows representative image of the distribution of cAMP signal of the cytosolic EPAC-S^H187^ FRET sensor Neonatal Mouse Atrial Myocytes (NMAMs). In WT NMAMs, in control conditions, the FRET change in response to 3 μM PE represented 13.1 ± 1.5% peak of the saturating response (Figure 5B). ST034307, an AC1 inhibitor, did not cause a significant reduction in response to 3 µM PE (11.3 ± 2.4 %, P=0.8583, Figure 5A). Bafilomycin A1 caused a reduction in response to PE to 4.6 ± 0.8% (n=7, N=4, P<0.001, Figure 5B). Addition of NAADP pathway inhibitors (1 µM of *trans*-Ned-19 and 10 µM of SG-094) both caused a significant reduction in PE responses compared to control (Figure 5B).

**Figure 5:**
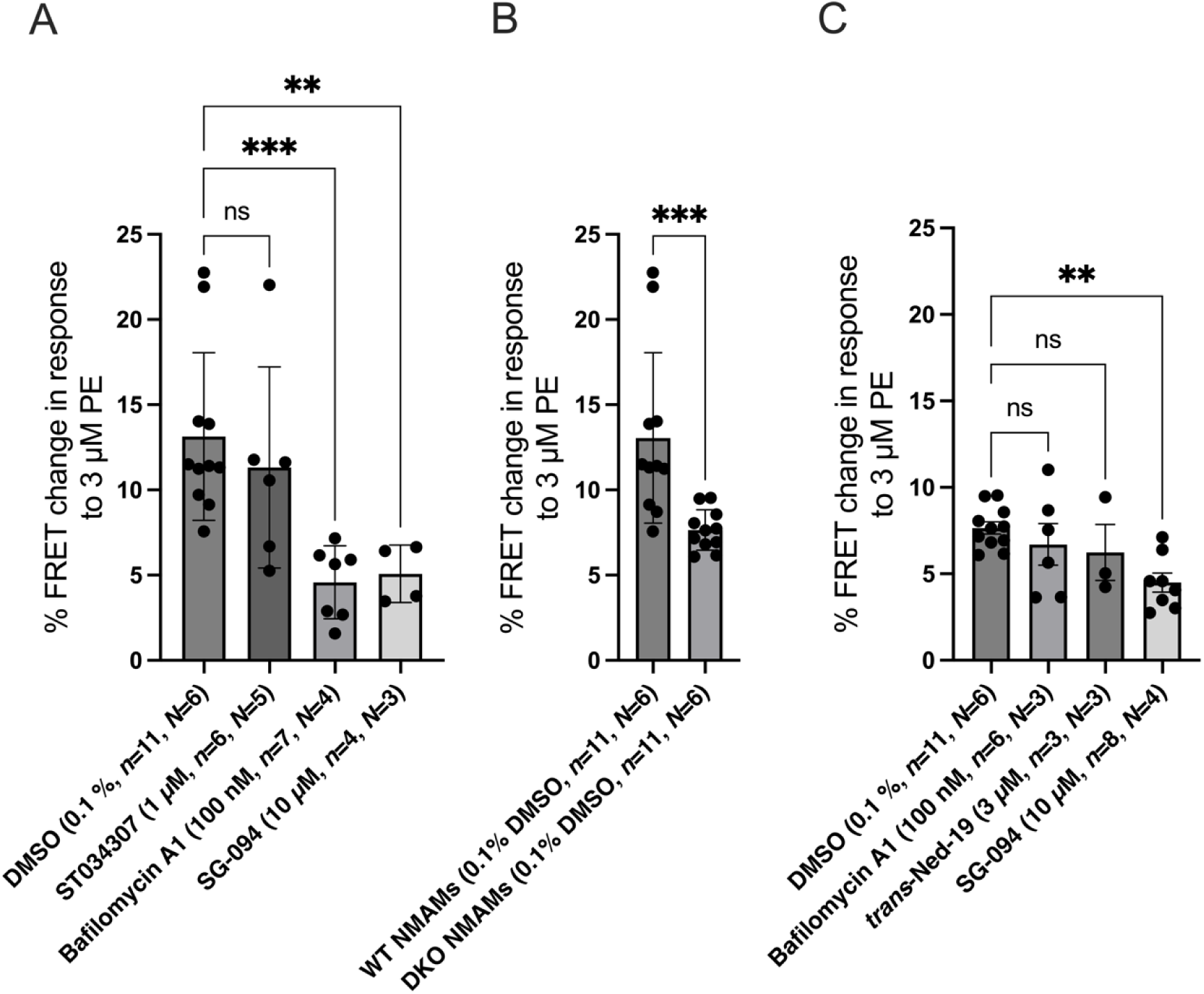
Acute α-adrenergic stimulation leads to lysosomal Ca^2+^ signalling via AC1 and AC8 activation. **A:** Quantification of the peak FRET change in response to 3 µM PE in WT NMAM in the presence of either 0.1 % DMSO (*n*=11, N=6), 1 µM ST034307 (*n*=6, N=5), 100 nM Bafilomycin A1 (*n*=7, N=4), 1 µM trans-Ned-19 (*n*=6, N=5) or 10 µM SG-094 (*n*=4, N=3) in E1 buffer. **B:** Quantification of the peak FRET change in response to 3 µM PE in WT NRAMs in the presence of DMSO (*n*=11, N=6) and in DKO NMAM in the presence of DMSO (*n*=11, N=6) in E1 buffer. **C**: Quantification of the peak FRET change in response to 3 µM PE in DKO NMAM in the presence of either 0.1 % DMSO (*n*=11, N=6), 100 nM Bafilomycin A1 (*n*=6, N=3), 1 µM trans-Ned-19 (*n*=3, N=3) or 10 µM SG-094 (*n*=8, N=4) in E1 buffer. Data are represented as mean ± SEM, panel A and B were analysed using Kruskal Wallis followed by Dunn’s multiple comparison and panel C a non-parametric Mann-Whitney U test; n=experiments; N=isolations; non-significant (ns)=P>0.05, **P<0.01, ***P<0.001.

In contrast, activation of α-adrenergic pathway using PE resulted in a FRET change of 7.7 ± 0.3 % in DKO NMAMs. This is a significant difference compared to the responses seen in WT NMAMs (P<0.01, Figure 5C). Addition of Bafilomycin A1 or *trans*-Ned-19 to DKO NMAMs did not cause significant reduction in response to 3µM PE, unlike in NRAMs. SG-094 (10 µM) caused a reduction in peak response to PE compared to DKO control (4.5 ± 0.5 %, n=8, N=4, P<0.01, Figure 5D).

### [5] AC1 and AC8 contribute to Ca^2+^ transient amplitude in response to α-adrenergic stimulation

Adult WT and DKO isolated atrial myocytes were loaded with the cell-permeant Ca^2+^ sensitive dye Fluo-5F-AM to measure changes in cytosolic Ca^2+^ in response to PE (Figure 6Ai). Normalised Ca^2+^ transient amplitude is expressed as change in mean cell fluorescence (F-F0)/F0 in the cell (Figure 6B). When stimulated at 1 Hz in Tyrode solution at 37°C, atrial myocytes exhibited the classical pattern of Ca^2+^ transient (Figure 6Aii-iii). Basal Ca^2+^ showed no significant differences between both groups. As shown in Figure 6B, knockout of AC1 and AC8 did not alter the basal Ca^2+^ compared to WT atrial myocytes. Perfusion with 10 µM PE resulted in an increase in Ca^2+^ transient amplitude by 26.6 ± 3.9 % in WT atrial myocytes at 1 min (n=10, N=3, Figure 6C and 6D). DKO atrial cells showed an increase in Ca^2+^ transient amplitude of 1.1 ± 4.4 % (n=10, N=3), a significantly attenuated response to PE compared to WT. Responses to PE in both WT and DKO had a tendency to decline over the total exposure time to PE (Figure 6C and 6D).

**Figure 6:**
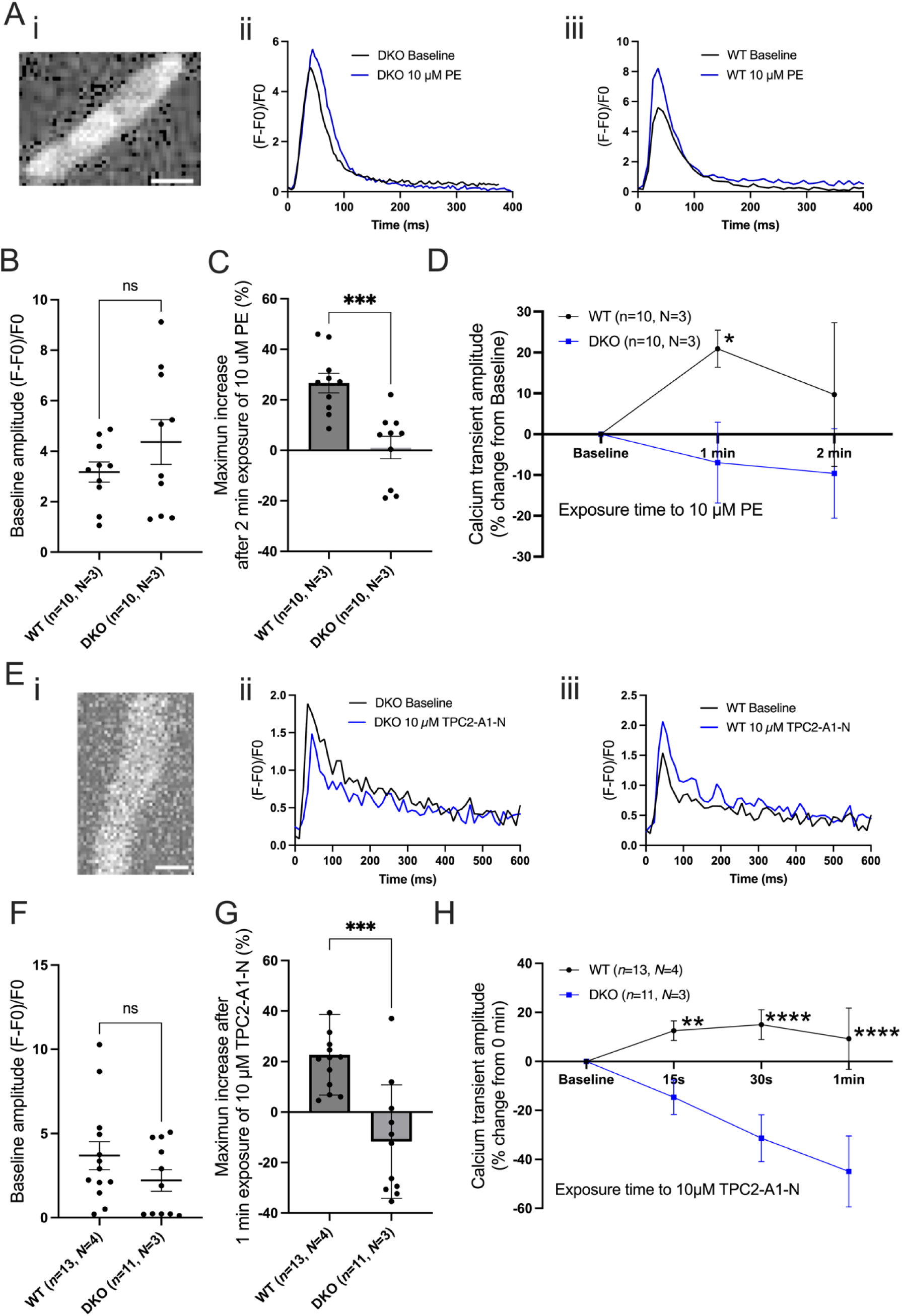
Expression of AC1 and AC8 contributes to Ca^2+^ transient amplitude in response to α-adrenergic stimulation and TPC2 stimulation. **Ai**: Representative image of a DKO arial myocyte loaded with 3 µM Fluo-5F. **ii**: Representative traces to show spontaneously generated Ca^2+^ transients in isolated atrial myocytes WT C57s in control condition (black) and after exposure to 10 µM PE (blue). **iii**, Representative traces to show spontaneously generated Ca^2+^ transients in isolated atrial myocytes DKO C57s in control condition (black) and after exposure to 10 µM PE (blue). **B**: Baseline Ca^2+^ transient amplitude in isolated WT (n=10, N=3) and DKO (n=10, N=3) C57s atrial myocytes in control conditions in Tyrode. **C**: Maximum increase (in %) recorded over 2 minutes of exposure to 10 µM PE in isolated WT (n=10, N=3) and DKO (n=10, N=3) C57s murine atrial myocytes. **D**: Ca^2+^ transient amplitude response (in %) in isolated WT (n=10, N=3) and DKO (n=10, N=3) C57s murine atrial myocytes after exposure of 10 µM PE over 2 minutes. **Ei**: Representative image of a DKO arial myocyte loaded with 3 µM Fluo-5F. **ii**: Representative traces to show spontaneously generated Ca^2+^ transients in isolated atrial myocyte WT C57s in control condition (black) and after exposure to 10 µM TPC2-A1-N (blue). **iii**: Representative traces to show spontaneously generated Ca^2+^ transients in isolated atrial myocyte DKO C57s in control condition (black) and after exposure to 10 µM TPC2-A1-N (blue). **F**: Baseline Ca^2+^ transient amplitude in isolated WT (n=10, N=3) and DKO (n=10, N=3) C57s atrial myocytes in control conditions in Tyrode. **G**: Maximum increase (in %) recorded over 2 minutes of exposure to 10 µM TPC2-A1-N in isolated WT (n=10, N=3) and DKO (n=10, N=3) C57s murine atrial myocytes. **H:** Ca^2+^ transient amplitude response (in %) in isolated WT (n=10, N=3) and DKO (n=10, N=3) C57s murine atrial myocytes after exposure of 10 µM TPC2-A1-N over 1 minute. Data are represented as mean ± SEM and were analysed using paired t-test in panel A and using 2-way repeated measures Scale bar represents 20 μm in panel Ai and Ei. ANOVA followed by Šídák’s multiple comparisons test in panel B; n=experiments; N=animals; *P<0.05, **P<0.01, ***P<0.001, ****P<0.0001.

To investigate the role of NAADP pathway activation in WT and DKO mice, changes in cytosolic Ca^2+^ transients were measured in response to TPC2-A1-N, a TPC2 activator (Gerndt, Chen et al. 2020). When stimulated at 1 Hz in Tyrode at 37°C, atrial myocytes also exhibited the classical pattern of Ca^2+^ transient in response to 10 µM TPC2-A1-N (Figure 6Eii-iii). DKO cells (2.2 ± 0.8, n=13, N=4) did not alter the basal Ca^2+^ compared to WT (3.7 ± 0.6, n=11, N=3) atrial myocytes (Figure 6F). Addition of 10 µM of TPC2-A1-N to the perfusion solution resulted in an increase in Ca^2+^ transients in WT and not in DKO (Figure 6G and 6H). Maximum percentage increase in Ca^2+^ transient amplitude was measured after 1 min exposure of TPC2-A1-N. In the WT we observe an increase in amplitude over the first 30 s and then atrial myocytes had a tendency to decline during TPC2-A1-N exposure time frame. DKO amplitude decreases over 1 min of exposure (Figure 6H). In WT cells, TPC2-A1-N caused the Ca^2+^ transient amplitude to increase by 22.7 ± 4.4 % (n=11, N=3) while in DKO atrial cells, TPC2-A1-N resulted in a fall in Ca^2+^ transient amplitude of 11.7 ± 6.8 % (n=13, N=4), representing a significant difference in response to TPC2 activation (P<0.001, Figure 6G and 6H).

### [6] Lysosomal Ca^2+^ signalling pathway contributes to the increase in cellular cAMP in human iPSC derived atrial myocytes

We used specific atrial-like cardiomyocytes derived from human-induced pluripotent stem cells (hiPSC-aCMs) to investigate the effect of the TPC2 agonist TPC2-A1-N on cAMP production. We labelled the live cells with lysotracker to confirm the presence of lysosomes (Figure 7A). FRET experiments were carried out on hiPSC-aCMs to build on the FRET data collected in NRAMs and to demonstrate the translational value of these human cells to study the NAADP pathway. Figure 7B presents cAMP levels in response to 10 µM TPC2-A1-N in the presence or absence of TPC2 antagonist 10 µM SG-094. Peak cAMP levels were significantly lower in the presence of SG-094 with a FRET change of 22.6 ± 6.2 % (n=6, N=3) in response to 10 µM TPC2-A1-N compared to controls (34.5 ± 4.1 %, n=4, N=3, P<0.05, Figure 7B).

**Figure 7.**
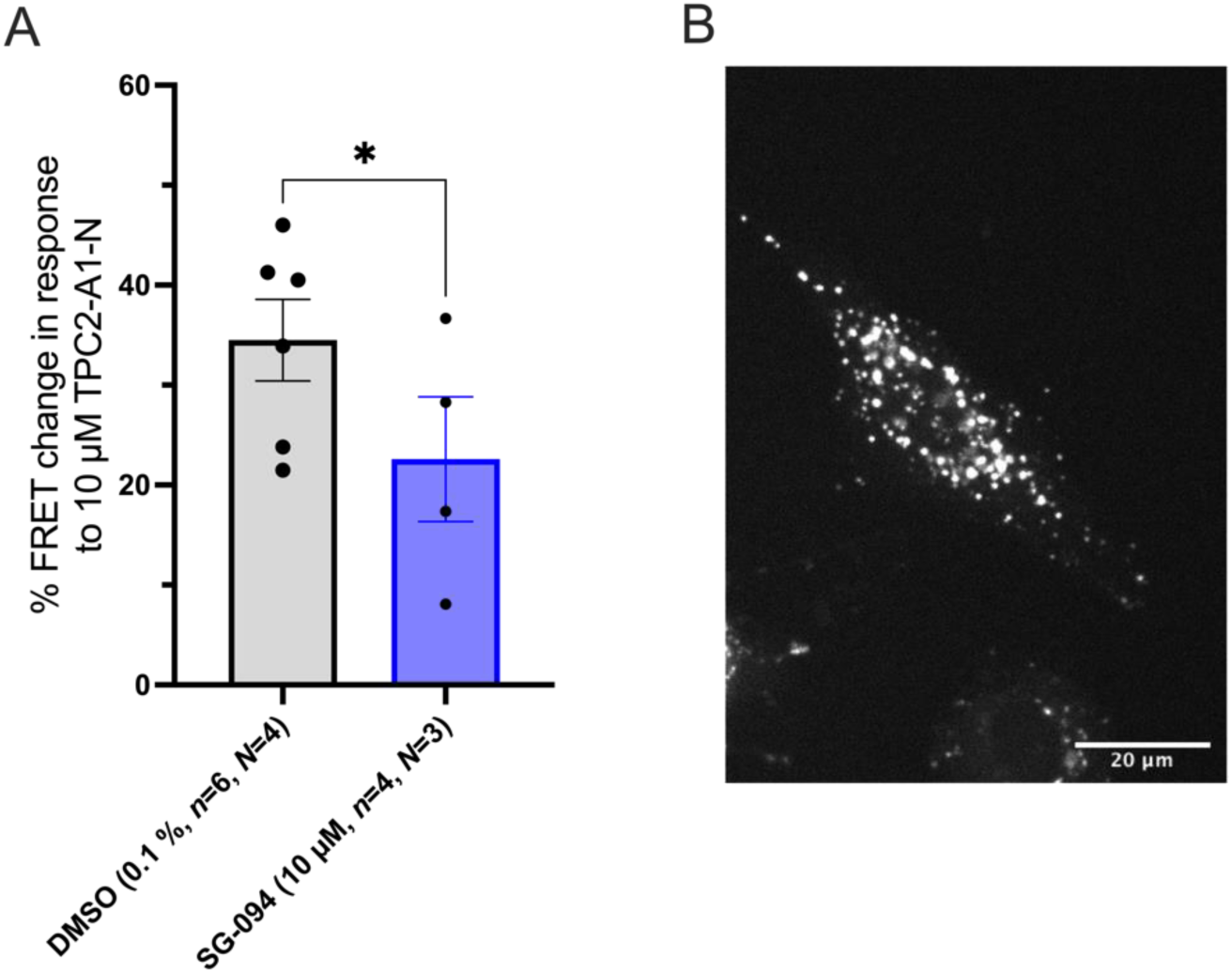
Providing first evidence of NAADP pathway in human iPSC-aCMs. **A:** Representative image of iPSC-aCMs loaded Lysotracker **B:** Quantification of the peak FRET change in response to 3 µM PE in iPSC-aCMs in the presence of either 0.1 % DMSO (*n*=6, N=4) or 10 µM SG-094 (*n*=4, N=3) in E1 buffer. Scale bar in B represent 20 µm. Data are represented as mean ± SEM and were analysed using paired t-test comparison. Normality was assessed using the Shapiro-Wilk test; n=experiments; N=isolations; * P<0.05.

## DISCUSSION

This manuscript focuses on NAADP signalling in the cardiac atrium in response to α-adrenergic stimulation and presents the novel finding that NAADP mediated Ca^2+^ release from lysosomes, and IP_3_ mediated Ca^2+^ release impacts atrial function. We investigated whether downstream activation of Ca^2+^ sensitive ACs (AC1/8) and release of lysosomal Ca^2+^ in response to IP_3_R activation, via an α-adrenoceptor-linked signalling pathway, contributes to atrial myocyte function. In whole tissue, pharmacological inhibition of the NAADP pathway reduced the contractile response to PE, our functional work showed that inhibition of NAADP signalling by addition of 500 μM BZ-194, 1 µM *trans*-Ned-19 or 100 nM Bafilomycin A1 significantly reduced the positive chronotropic effect of PE (Figure 1). Additionally, Knockout of AC1 and AC8 also significantly reduced the positive chronotropic effect of PE (Figure 3). These datasets indicate the downstream effects of α-adrenergic signalling are dependent upon NAADP mediated Ca^2+^ signalling from lysosomes in chronotropy and inotropy in SAN and atrial myocytes. Furthermore, FRET experiments show a decrease in cAMP activity in response to 3 µM PE following addition of several NAADP inhibitors but is not seen in NRVMs. These results provide strong evidence that cAMP is reduced by NAADP pathway inhibitors, indicating a greater role for acidic Ca^2+^ stores in atrial function. We investigated DKO of Ca^2+^- sensitive AC1 and AC8 in mice as a possible explanation for these observations as lysosomal Ca^2+^ release could be activating AC1 and/or AC8 in α-adrenergic response. In intact DKO atria there was a reduction in positive chronotropy and inotropy observed in response to PE, and in adult isolated atrial myocytes the rise in Ca^2+^ transient amplitude was attenuated in response to PE and a TPC2 activator.

Data in Figure 5.A present the effects of Bafilomycin A1, *trans*-Ned-19 and SG-094 on lysosomal Ca^2+^ release in response to PE stimulation in WT NMAMs. In DKO NMAMs, the response to PE is reduced when compared to WT (Figure 5.B). Bafilomycin A1 and *trans*-Ned-19 show no effect on cAMP activity in response to PE in DKO NMAMs (Figure 5.C). SG-094 significantly reduces the response to PE in the DKO NMAMs similar to WT NMAMs. We observed a similar reduction in the cAMP rise in response to PE in WT NMAMs in the presence of inhibitors (*trans*-Ned-19, Bafilomycin A1 and SG-094) and in control condition (DMSO) DKO NMAMs (Figure 5.B). The FRET finding that Bafilomycin A1 has no further effect on cAMP activity following DAC1/8 KO suggests the effect of Bafilomycin A1 in response to PE requires the presence of AC1/8 for a cAMP response, and this can be explained by AC1/8 activity being downstream of the lysosomes Ca^2+^ handling upon α-adrenergic stimulation.

ST034307, an AC1 inhibitor, caused a non-significant reduction in cAMP activity in response to 3 µM PE (11.3 ± 2.4 %, Figure 5.A). The Hom AC1 Het AC8 genotype cause a significant reduction (similar to DKO) of maximum rate change of spontaneously beating mouse right atrial tissue exposed to PE compared to WT whereas Het AC1/8 and Het AC1 Hom AC8 caused a non-significant reduction (Supplementary Figure 3.C). Previous work, showed ST034307 to inhibit rate change of spontaneously beating mouse right atrial tissue but not tension generated in response to electrical stimulation mouse left atrial tissue during PE exposure and reduced the beating rate of isolated guinea pig atrial and SAN cells loaded with Fluo-5F-AM exposed to PE but no reduction in the amplitude of the CaT (Bose, Read et al. 2022). AC1 appears to play a greater role in rate regulation in the SAN and contributes less to inotropic responses during α-adrenergic stimulation. No localisation studies of TPC2 or AC8 were able to be performed. However, immuno-fluorescent labelling of isolated guinea pig SAN myocytes with LAMP2 and AC1 were conducted and presented colocalization, pixel by pixel analysis presented a Pearson’s colocalisation overlap coefficient of R=0.8095 ± 0.04565 (Capel, Akerman et al. 2024) and AC1 and IP_3_R2 receptors have been identified to be located in close proximity in atrial myocytes (Capel, Bose et al. 2021) and in SAN myocytes (Bose, Read et al. 2022).

Recently, Yuan et al investigated the interplay between the lysosomal TPC2 channels and endoplasmic reticulum (ER)-localized IP_3_Rs, providing experimental evidence that TPC2-mediated local Ca^2+^ release gets amplified by Ca^2+^ depletion from the ER via IP_3_Rs to give rise to global Ca^2+^ signals (Yuan, Arige et al. 2024). Taken together, the PE dose response results (reduction of response in the presence of NAADP pathway inhibitors), the FRET results (reduced cAMP activity in the presence of NAADP pathway inhibitors), the atrial Ca^2+^ transient results (reduction of response to PE and TPC2-A1-N in DKO) and the DKO experiments (abolish of reduction seen in WT or control conditions), support a potential for cross talk. We show the involvement of AC1/8 and lysosomal Ca2+ signalling in the cAMP response to PE involving the interaction between AC1/8 and lysosomal Ca2+ signalling, an illustration of the proposed model is presented in Figure 8.

**Figure 8:**
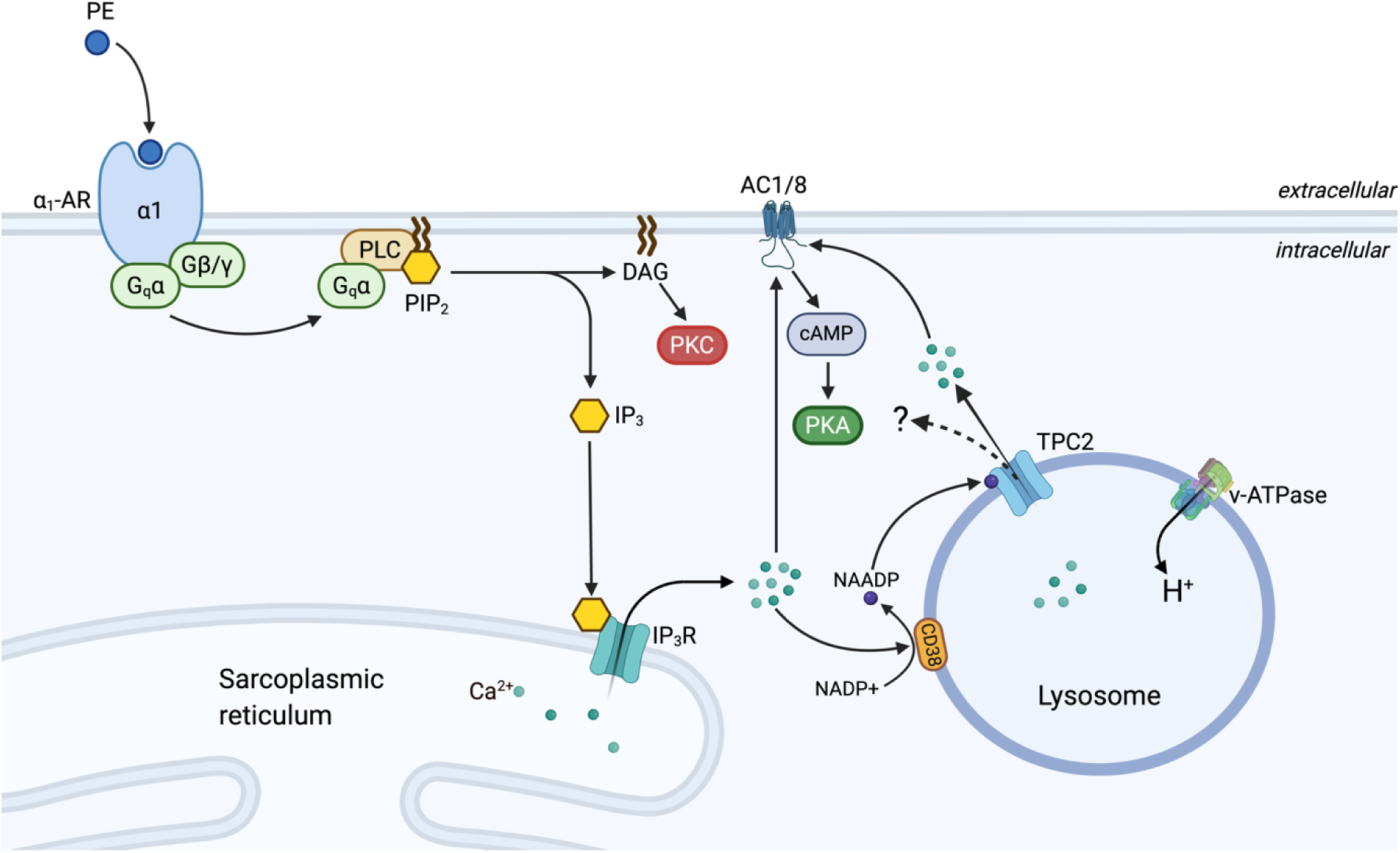
Proposed simplified model of crosstalk between IP_3_ and NAADP. Scheme whereby crosstalk between IP_3_ and NAADP mediated Ca^2+^ release regulate the release of Ca^2+^ from intracellular stores in atrial myocytes and influence atrial cardiac function and pacemaking. Activation of α1-AR leads to elevated IP_3_; IP_3_ activates IP_3_R resulting in release of Ca^2+^ from the SR, which subsequently leads to activation of AC1 and/or AC8, and causes elevation of NAADP levels. NAADP acts to release Ca^2+^ from lysosomes through TPC2 channels also leading to activation of AC1 and/or AC8. Activation of α1-AR (by PE) leads to increased atrial cytoplasmic Ca^2+^ transients and cAMP production. The location of CD38 is not yet established and may perhaps be associated with the SR (Lin, Bolton et al. 2017). The abbreviations used are as follows: α-AR: Alpha-adrenoreceptor; PLC: Phospholipase C; DAG: Diacylglycerol; PIP2: Phosphatidylinositol 4,5-bisphosphate, PKC: Protein kinase C. Created with BioRender.com.

Studies from over a decade ago uncovered β-adrenergic receptor signalling increases NAADP levels in whole heart (Lewis, Aley et al. 2012, Nebel, Schwoerer et al. 2013). Nebel et al. reported a role for NAADP in arrhythmogenic Ca^2+^ release in cardiomyocytes evoked by β-adrenergic stimulation (Nebel, Schwoerer et al. 2013). In 2011, Collins et al. published data showing NAADP influences excitation-contraction coupling by releasing Ca^2+^ from lysosomes in atrial myocytes using photorelease methods of a caged NAADP compound (Collins, Bayliss et al. 2011). In the same study, isoprenaline (4 nM) stimulated Ca^2+^ release from lysosomes providing evidence for a role of NAADP in regulating Ca^2+^ signalling in guinea-pig atrial myocytes.

It has also been shown that the involvement of IP_3_R in response to extracellular stimuli redistributes Ca^2+^ from the ER to the cytosol and also allows extracellular stimuli to regulate Ca^2+^ exchange between the ER and lysosomes by directly releasing lysosomal Ca^2+^ to IP_3_R on the ER (Atakpa, Thillaiappan et al. 2018). The study by Yuan et al mentioned earlier, in HeLa and HEK cells revealed that TPC2 opening is coupled to IP_3_Rs and that Ca^2+^ signals originating from NAADP-sensitive channels on lysosomes exhibit a major dependence on ER Ca^2+^ despite their lysosomal origin (Yuan, Arige et al. 2024). Such a mechanism has not yet been observed in cardiomyocytes. However, experiments using isolated cardiomyocytes from TPC1/2 knockout mice demonstrated reduction in contractility and decrease Ca^2+^ content in the SR (de Zélicourt, Fayssoil et al. 2024).

The present study is the first time the AC1/8 DKO colony has been used to investigate cardiac physiology and pharmacology. We used HREM methods (Weninger and Mohun 2007) for visualisation of DKO hearts and compared them to WT to confirm any anatomical-pathological changes due to the knockout of AC1/AC8 genes; Analysis of tissue architecture and morphology showed no phenotypic differences in any chamber between WT and DKO hearts that might suggest possible compensatory mechanisms. There was a significant difference seen in the change in spontaneous beating rate following exposure to metoprolol between WT and DKO right atria. WT right atria demonstrated a larger reduction in spontaneous beating rate compared to DKO. Different Het and Hom genotypes for AC1 and AC8 that were generated unravelled the extent to which AC1 or AC8 were implicated in the response to PE, the Hom for AC1/Het for AC8 genotype showed a significant increase in EC_50_ of 3.7 (3.2–9.2) μM compared to WT and DKO, highlighting the implication of AC1 in rate response. This finding raises questions regarding the compensatory mechanisms relating to adaptation to AC1/AC8 DKO. Possible mechanisms include modulation of Ca^2+^ signalling pathways within the cell such as upregulation SR channels (IP_3_Rs or RyRs). Ca^2+^ transient investigations using direct stimulation of TPC2 channel opening (ligand biased towards TPC2 Ca^2+^ flux, 10 μM TPC2-A1-N or 3 µM PE caused a Ca^2+^ transient amplitude increase in WT atrial myocytes, similar to previous work by Collins et al, using caged NAADP (Collins, Bayliss et al. 2011). Experiments repeated in DKO atrial myocytes did not respond to 10 µM TPC2-A1-N or showed a decrease over exposure time in the presence of 3 µM PE. Ca^2+^ transient work presented The study reveals a link between lysosomal Ca^2+^ release and direct stimulation of AC1 and/or AC8. As the stimulation of the NAADP pathway or α-adrenergic pathway in DKO completely abolished the increase in Ca^2+^ transient and even presented a decrease of amplitude for the total duration of the protocol, underlining a crucial role for NAADP signalling and highlighting AC1 and/or AC8 as key activators/drivers in Ca^2+^ from the lysosomes.

FRET experiments conducted using hiPSC-aCMs confirmed that activation of the NAADP pathway via TPC2-A1-N leads to the generation of cAMP (Figure 7). These results highlight the relevance of these findings in human models of cardiac physiology. Disturbances of cardiac atrial rhythm lead to inefficient ventricular filling and low flow states that may lead to reduced cardiac output and arterial thrombotic complications such as stroke. The most common sustained atrial arrhythmia is AF. The prevalence of AF continues to increase as the population ages (Workman, Kane et al. 2008, Samuthpongtorn, Jereerat et al. 2021). Lysosomal dysfunction has been characterised in several cardiac conditions (Chi, Riching et al. 2020), including congenital atrial septal defects where lysosomes in the right atrium are increased in patients (Kottmeier and Wheat 1967). Further progressive changes, such as the accumulation of lysosomes, have been found to correlate with atrial cellular electrophysiological changes (Mary-Rabine, Albert et al. 1983). These findings support a role for lysosomes as an important intracellular organelle in cardiovascular disease and targeting lysosomes and NAADP pathway maybe of therapeutic interest in these patients by targeting this signalling pathway specific to atrial tissue.

IP_3_R (Berridge 2009) and AC1 and AC8 (Mattick, Parrington et al. 2007) being linked to pacemaking (Bose, Read et al. 2022) and unravelling the interaction between IP_3_R pathways and ACs provides important insights into mechanisms for initiation of atrial arrhythmias. FRET studies on hiPSC-aCMs confirms the presence and significance of the NAADP pathway in these human cardiac cell models and warrants further investigations of NAADP signalling in atrial pathophysiology.

## Conclusion

In summary, this study brings together two major aspects of cardiac atrial Ca^2+^ handling: IP_3_-mediated Ca^2+^ release from the SR and Ca^2+^ release from lysosomes via NAADP signalling. We have demonstrated that the NAADP pathway is involved in the α-adrenergic response, highlighting that lysosomal Ca^2+^ is an important component of α-adrenergic stimulation in the cardiac atria acting on AC1/8 (Figure 8). Lysosomal Ca^2+^ signalling contributes to cellular cAMP generation in the cardiac atria and presents an important pathway for the control of sino-atrial node activity and atrial inotropy, whether Ca^2+^ release via TPC2 acts on other targets to regulate atria function remains to be investigated (Figure 8). Derangement of this signalling pathway can possibly contribute to cardiac pathologies such as AF. The observations we present in this study strengthens the relevancy of the NAADP pathway acting via lysosomal Ca^2+^ in atrial cardiomyocytes.

## Supplement Figures and Legends

**Supplementary Figure 1:**
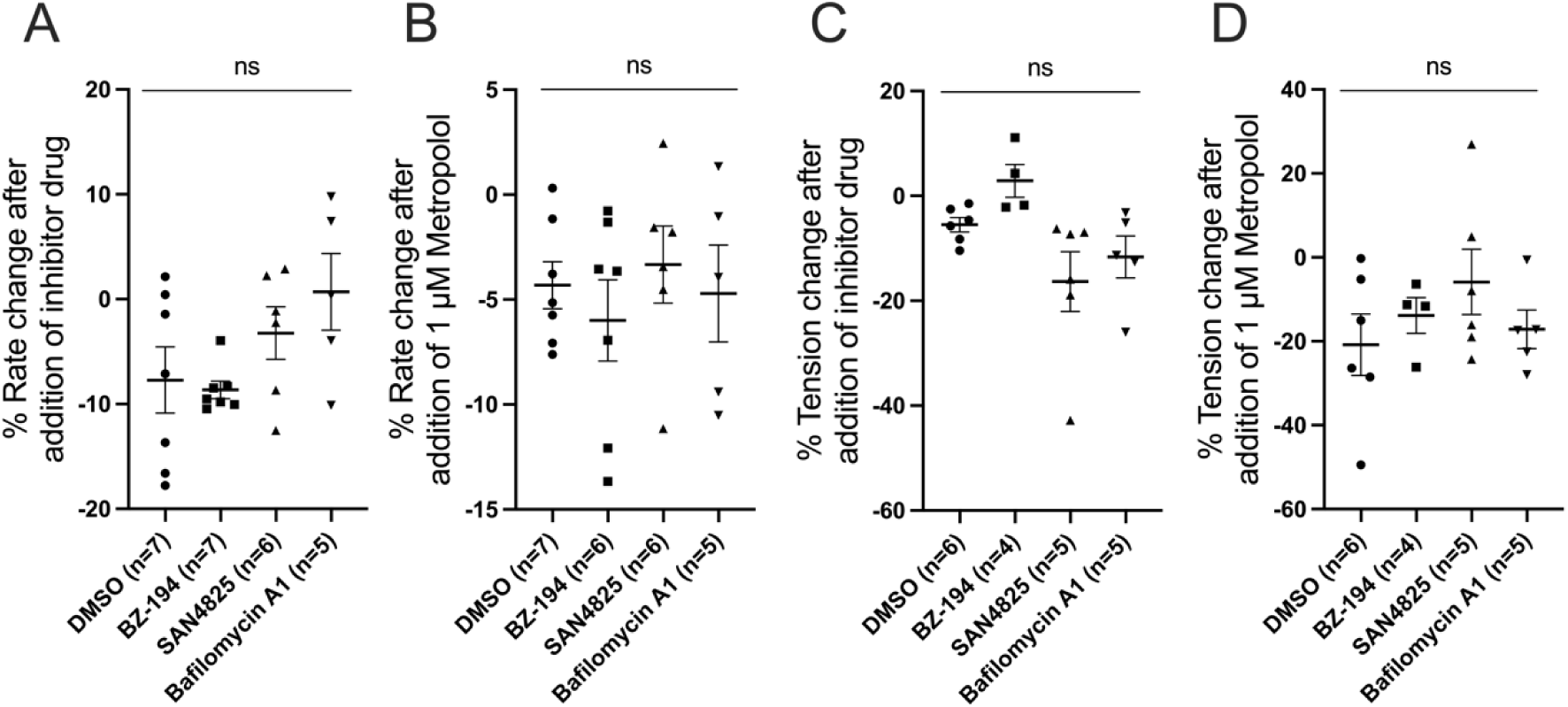
Baseline beating rate and tension comparison shows no significant differences. **A**: Comparison of percentage decrease rate after addition of 0.001 % DMSO, 500 μM BZ-194, 10 μM SAN4825 or 100 nM Bafilomycin A1. **B**: Comparison of percentage decrease rate after addition of 1 µM Metoprolol, under control conditions (0.001 % DMSO) or in the presence of 500 μM BZ-194, 10 μM SAN4825 and 100 nM Bafilomycin A1. **C**: Comparison of percentage decrease in tension after addition of 0.001 % DMSO, 500 μM BZ-194, 10 μM SAN4825 or 100 nM Bafilomycin A1. **D**: Comparison of percentage decrease in tension after addition of 1 µM Metoprolol, under control conditions (0.001 % DMSO) or in the presence of 500 μM BZ-194, 10 μM SAN4825 and 100 nM Bafilomycin A1. Normality was assessed using the Shapiro-Wilk test. Data were analysed using 2-way repeated measures ANOVA followed by Dunnett’s multiple comparisons test. non-significant (ns)=P>0.05.

**Supplementary Figure 2:**
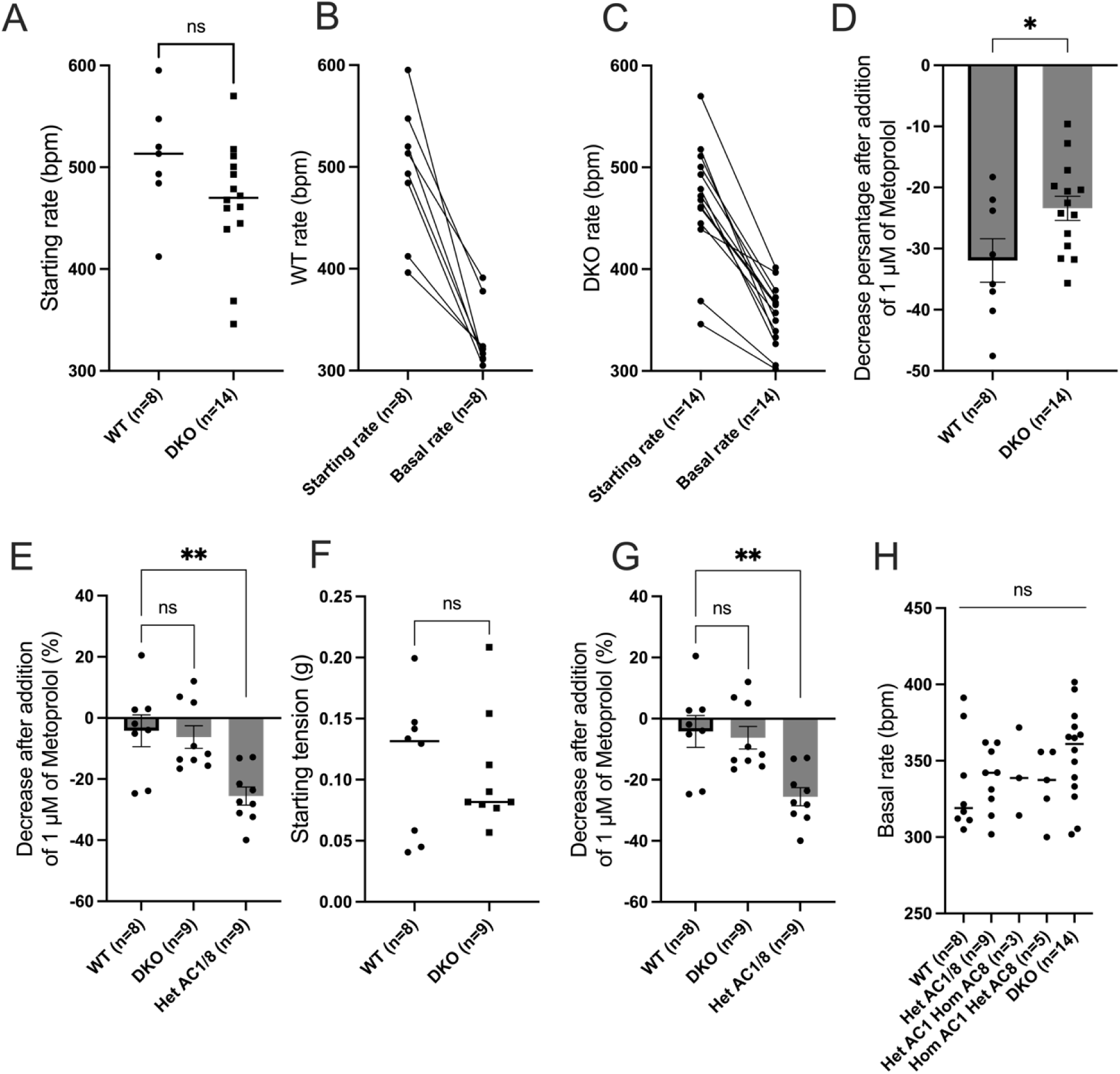
No significant changes observed in baseline beating rate and tension between WT and DKO right and left atrial preparation before addition of PE. **A**: Comparison of spontaneously starting beating rate of WT (n=8) and DKO (n=14) C57s murine right atrial preparations in PSS prior to addition of drug or stimulation with PE. **B**: Comparison of spontaneously starting and basal beating rate of WT (n=8) C57s murine right atrial preparations in PSS. **C**: Comparison of spontaneously starting and basal beating rate of DKO (n=14) C57s murine right atrial preparations in PSS. **D**: Comparison of percentage decrease rate between starting rate and basal rate of WT (n=8) and DKO (n=14) C57s murine right atrial preparations. **E**: Comparison of starting tension of WT (n=8) and DKO (n=10) C57s murine right atrial preparations in PSS prior to addition of drug or stimulation with PE. **F**: Comparison of percentage decrease contractile force between starting rate and basal rate of WT (n=8) Het AC1/8 (n=7) and DKO (n=9) C57s murine right atrial preparations. **G**: Comparison of percentage decrease rate between starting rate and basal rate of WT (n=8) and DKO (n=9) C57s murine right atrial preparations. **H**: Comparison of spontaneously baseline tension WT (n=8), Het AC1/8 (n=9), Het AC1/HomAC8 (n=3) and HomAC1/HetAC8 (n=5) and DKO AC1/8 (n=14) C57s murine right atrial preparations in PSS after addition of 1 µM Metoprolol. Data are represented as mean ± SEM and were analysed using paired t-test comparison or 2-way repeated measures ANOVA followed by Dunnett’s multiple comparisons test; * P<0.05, ** P<0.01.

**Supplementary Figure 3:**
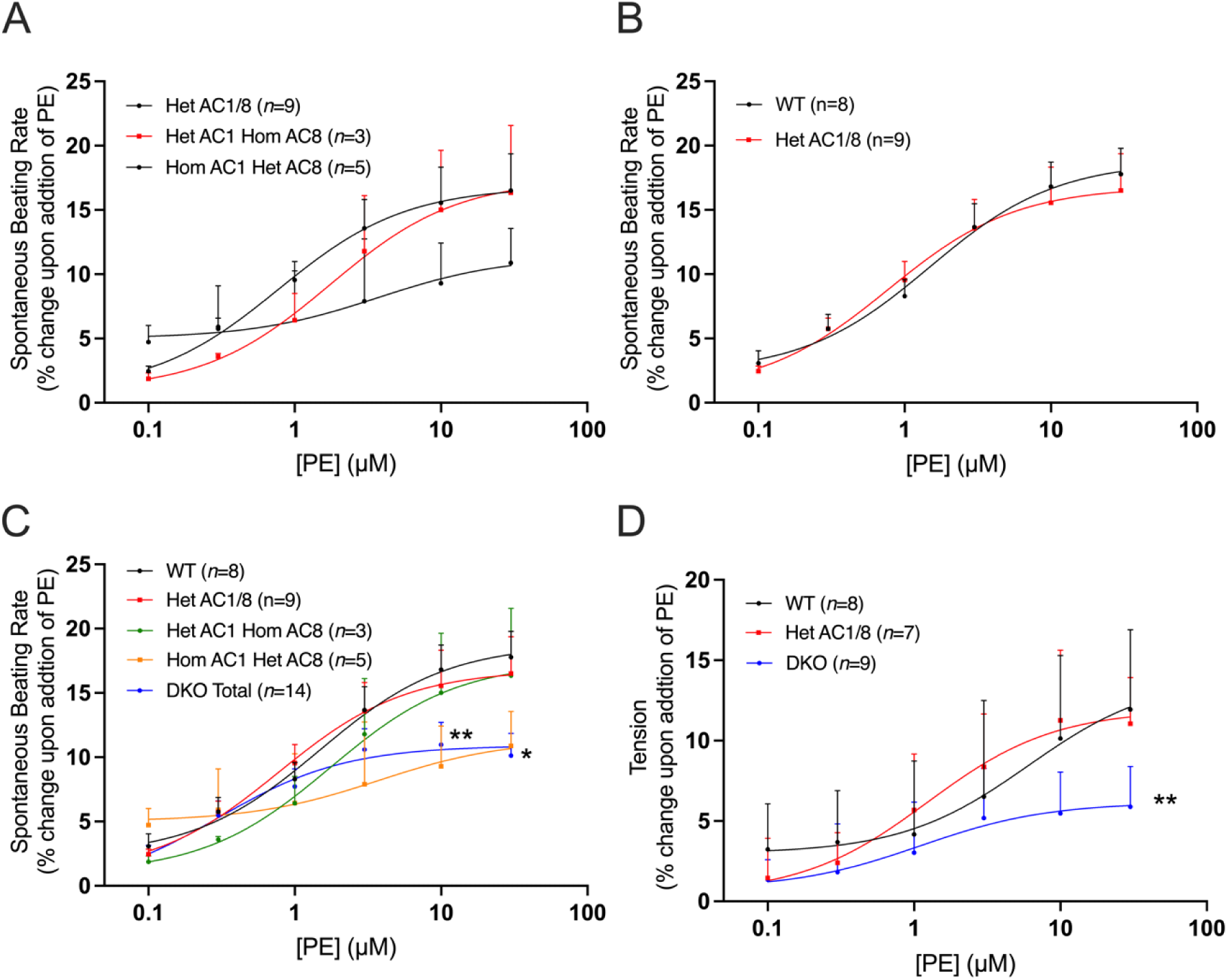
Dose-response curves observed in beating rate and tension between different mouse genotypes (AC1/AC8) in right and left atrial preparation in response to PE. **A**: Rate responses to PE (0.1 µM - 30 µM) in spontaneously beating murine right atrial preparations in Het AC1/8 (n=5, red), Het AC1/HomAC8 (n=3, green) and HomAC1/HetAC8 (n=6, orange). **B**: Rate responses to PE (0.1 µM - 30 µM) in spontaneously beating murine right atrial preparations in WT (n=8, black) and Het AC1/8 (n=5, red). **C**: Rate responses to PE (0.1 µM - 30 µM) in spontaneously beating murine right atrial preparations in WT (n=8, black), Het AC1/8 (n=5, red), Het AC1/HomAC8 (n=3, green), HomAC1/HetAC8 (n=6, orange) and DKO AC1/8 (n=14, blue). **D**: Tension change to PE (0.1µM - 30 µM) in stimulated beating murine left atrial preparations in WT (n=8), HetAC1/8 (n=10) and DKO AC1/8 (n=9). Dose-response curves were fitted using log(agonist) vs. response (three-parameter model). Asterisks indicate significance level for effect of DKO compared to WT at individual concentrations as determined using 2-way repeated measures ANOVA followed by Šídák’s multiple comparisons test.

**Supplementary Figure 4:**
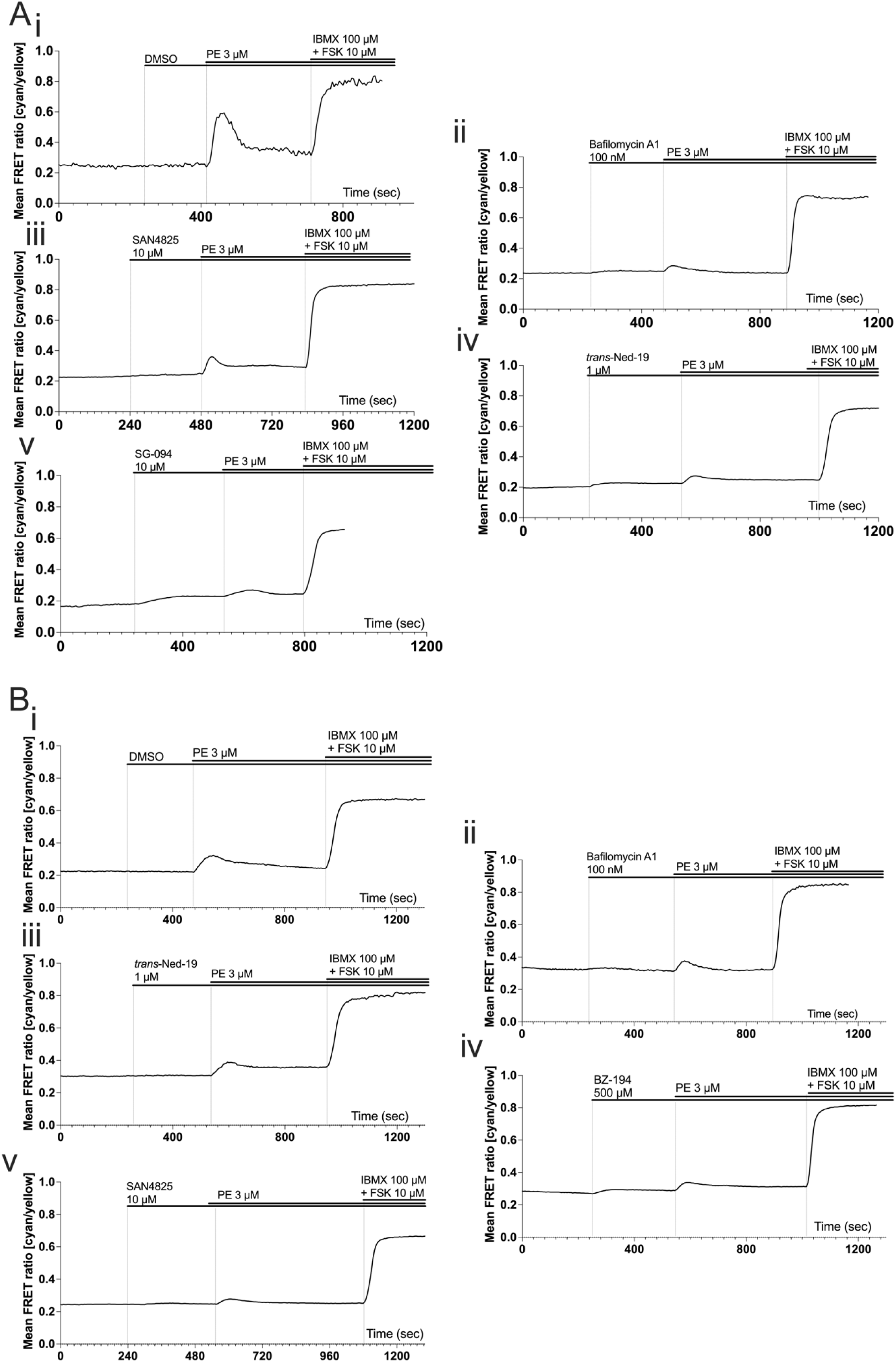
NRAMs and NRVMs Representative FRET traces in response to PE in the presence of NAADP inhibitors. **A**: Representative traces showing FRET change in single NRAMs expressing EPAC-S^H187^ in response to 3 µM PE under control conditions and following addition of 0.1 % DMSO (**i**), 100 nM Bafilomycin A1 (**ii**), 10 µM SAN4825 (**iii**), 10 µM *trans*-Ned-19 (**iv**) and 10 µM SG-094 (**v**). **B**: Representative traces showing FRET change in single NRVMs expressing EPAC-S^H187^ in response to 3 µM PE under control conditions (**i**) and following addition of 100 nM Bafilomycin A1 (**ii**), 1 µM Ned-19 (**iii**), 500 µM BZ-194 (**iv**) and 10 µM SAN4825 (**v**).

## Acknowledgments

This work was funded by the British Heart Foundation (BHF) project grant number PG/18/4/33521 (RABB and MZ). RABB was funded by a Sir Henry Dale Wellcome Trust and Royal Society Fellowship (109371/Z/15/Z) and acknowledges support from the Ellis T Davies Fellowship Endowment, University of Liverpool. SJB was funded by the British Heart Foundation (PG/18/4/33521). RAC was funded by the Wellcome Trust and Royal Society. AG is a Wellcome Trust Senior Investigator and a Principal Investigator of the British Heart Foundation Centre of Research Excellence at the University of Oxford. LB and DBS was funded by the British Heart Foundation (FS/SBSRF/22/31022 and FS/17/55/33100), the Oxford BHF Centre of Research Excellence (RE/13/1/30181), and Additional Ventures. MZ is supported by the British Heart Foundation (RG/17/6/32944). BVLP is a Wellcome Trust Senior Investigator (grant 101010). We thank Prof Patrizia Dell’Era, Brescia Italy (now deceased) for technical hiPSC-CM support. For the purpose of Open Access, the author has applied for CC BY public copyright licence to the any Author Accepted Manuscript version arising from this submission.

## Competing Interests

None to declare.

## Author contributions

RABB designed and lead the study. MZ designed, oversaw all the FRET studies. RABB, DAT, RC, EA formulated hypotheses. RABB, DAT, AG, MZ and EA contributed intellectually to the project. RABB and EA made the first draft. EA, RC, SB performed the experiments. EA performed statistical analysis. AK, MR, SB and MZ contributed to the FRET design and experimentation. SC contributed to hiPSC experiments. MK, FB and BVLP contributed to drugs developed and utilised in this study. QS, DBS and LB conducted the 3D histology studies. AC, SC Lab created the double knockout mouse colony in the US and exported the mice to UK. EA created the figures. All authors contributed to writing of the paper.

## hiPSC ethics

Full consent was provided for hiPSC cells used.

## STAR Methods

## EXPERIMENTAL MODEL AND SUBJECT DETAILS

### Animals

All work detailed complies with the “Principles and standards for reporting animal experiments” (Grundy, 2015) and the Animal (Scientific Procedures) Act 1986. All experimental protocols (Schedule 1) were approved by the University of Oxford, Procedures Establishment License Number XEC303F12. All animals were housed maintained in a 12-h light/dark cycle with ad libitum access to standard diet and sterilized water.

Disruption of the AC1 gene in mice was done in 1995 by (Wu, Thomas et al. 1995) to generate KO of AC1 mice. KO AC8 mice were generated by (Muglia, Schaefer et al. 1999), by deletion of exon 1 and exon 2 of the 5’-flanking region of AC8 gene and disruption was confirmed by assay of AC8 activity. AC1 and AC8 homozygote (DKO) mice were mated to obtain doubly heterozygotic F1 progeny. F1 mice were then backcrossed to AC1 homozygous mice to afford an F2 progeny with a 1:1 (heterozygote:homozygote) distribution at the AC1 allele and a 1:1 (wild type:heterozygote) distribution at the AC8 allele. F2 mice heterozygous for the AC8 allele and homozygous for the AC1 allele were backcrossed to afford 25% doubly homozygous mice (DKO mice) (Wong, Athos et al. 1999). Males from this DKO mouse line was then sent to us by Conti Laboratory from Wayne State University, unfortunately once the animals reached Oxford, females were too old to be able to breed. We then generated DKO colony by backcrossing AC1/8DKO male mice with WT females and proceeded with same method. In the process of generating DKO mice, Het AC1/8, Het AC1/HomAC8 and HomAC1/HetAC8 mice were generated and used for left and right atrial preparations, an experimental method described below. The C57BL/6, CD1 mice and rat litters were obtained from Charles River (UK).

## METHOD DETAILS

### Animal Experiments

#### Immunostaining

Cardiomyocytes were fixed with 2% paraformaldehyde for 15 minutes. The cells were washed 3 times with PBS. Cells were then blocked with 10% Donkey serum, 0.3% BSA and 0.1% triton X100 in PBS for 1 hour. Following the removal of the blocking buffer, cells were incubated overnight in 4°C with primary antibodies (listed below). Cells were then washed 3 times with PBS and incubated in the dark at room temperature for 4 hours with the secondary antibodies (1:400) donkey anti-mouse IgG conjugated to Alexa Fluor 564 and (A10040, Invitrogen) goat anti-rabbit IgG conjugated to Alexa Fluor 488 (SA510166, Invitrogen). Cells were washed 3 times in PBS and coverslips were mounted on slides with Vectashield^TM^ with DAPI to stain the nuclei. The slides were viewed using a Nikon eclipse Ti inverted confocal microscope (Nikon) with a 63x/1.2 water objective Plan Apo VC 60xA WI DIC N2 lens. NIS-Element viewer (Nikon) was used to acquire multichannel fluorescence images. For detection of DAPI, fluorescence excitation was 350nm with absorption detected at 460nm. For detection of AlexaFluor 488, fluorescence excitation was at 488nm with emission collected 515nm. Excitation at 561nm was collected at 595nm for detection of AlexaFluor 568nm and imaged sequentially at 2048 × 2048 (12 bits).

#### Lysotracker

For the staining of acidic compartments (lysosomes) in live cardiac myocytes, LysoTracker^TM^ red was used from Thermo Fisher Scientific (L12492). LysoTracker^TM^ probes are fluorescent acidotrophic probes for labelling and tracking acidic organelles in live cells. LysoTracker^TM^ dyes consist of a fluorophore linked to a weak base that is only partially protonated at neutral pH, are freely permeant to cell membranes and typically concentrate in spherical organelles. To label atrial myocytes, 50nM LysoTracker^TM^ red was incubated with the cells at room temperature in the dark for 20 minutes in Tyrode (in mM: NaCl 136, KCl 5.4, NaHCO_3_ 12, sodium pyruvate 1, NaH_2_PO_4_ 1, MgCl_2_ 1, EGTA 0.04, glucose 5) to allow it to partition into the acidic organelles. Myocytes were then washed twice and left in Tyrode for live imaging using Nikon Eclipse Ti Inverted Fluorescence Microscope with a 40x oil-immersion objective with illumination light at 546 nm.

#### High resolution episcopic microscopy

High resolution episcopic microscopy (HREM) was conducted on murine WT and DKO hearts as previously described (Kalisch-Smith, Ved et al. 2021). Whole hearts were fixed overnight at room temperature using Bouin’s fixative (HT10132, Merck), then thoroughly rinsed in PBS and subjected to gradual dehydration in a methanol series, starting at 10% + 10% between each step, up to 90% and then 90 %, 95 %, 100 % for 2 hours each step on a rocker at room temperature. Hearts were then embedded using a JB-4 kit (Catalogue number 00226-1, Polysciences). Briefly, hearts underwent overnight infiltration at 4 °C with a 50:50 mixture of methanol and infiltration solution (JB-4 Solution A plus 1.25% (w/v) Benzoyl Peroxide Plasticizer); they were rinsed in 100% infiltration solution for 1 hour in the dark at room temperature on rocker to minimise residual methanol; and left to incubate for up to 1 week at 4 °C in fresh infiltration solution. Finally, 6% (v/v) JB-4 solution B was added to the infiltration solution to initiate polymerization. Subsequently, analysis was carried out using a 3D Optical high-resolution episcopic microscopy imaging system (Indigo Scientific). HREM data were evaluated using Horos 3.3.6 (https://horosproject.org), and Amira for Life & Biomedical Sciences version 2019.4 (Thermo Fisher Scientific).

#### Mouse Atrial preparations

Male mice (between 25-35g) were terminated by concussion followed by cervical dislocation, the chest cavity was opened, and the heart was rapidly excised and placed in warm, oxygenated physiological saline solution (PSS, in mM: NaCl 125, NaHCO_3_ 25, KCl 5.4, NaH_2_PO_4_ 1.2, MgCl_2_ 1, glucose 5.5, CaCl_2_ 1.8, pH to 7.4 with NaOH) oxygenated with 95% O_2_/5% CO_2_) containing heparin (10U/mL). Hearts were transferred to a dissection chamber under a microscope, ventricular tissue was discarded, and the atria were cleared of any overlying tissue such as cardiovascular tubing and adipose tissue. Right and left atria were separated, and loops of thin suture were tied to the lateral edges of each atrium by directly knotting around a small area of the tissue before being mounted to a force transducer in an organ bath. One loop was anchored to a hook, and the other tied to the tension transducer with a resting tension between 0.22-0.24g, a correct weight to visualise contraction without adding a strain to the atrium. The preparation was hung in an organ bath filled with PSS, maintained at 37°C and oxygenated with 95% O_2_/5% CO_2_. Tension data were digitised using a PowerLabs bridge amplifier and recorded on LabChart5 software (all from ADInstruments, UK). Chronotropic effect was measured from the time interval between contraction of spontaneously beating right atrium and beating rate was calculated in real-time from the upstroke of the tension signal using the Chart5 Ratemeter function. Inotropic effect was measured from stimulated left atrium at 5Hz during these experiments.

#### Phenylephrine dose response curves

After the frequency force of right atrial contractions and force of contraction of left atrial contractions had stabilised (variation of no more than 2 beats per minute (bpm) and 0.2g over a 10 minute period in the right and left atrium respectively), the effects of PE were investigated over a cumulatively increasing concentration range of 0.1 μM to 30 μM. Preparations were excluded if stabilised beating rate under control conditions only in the presence of PSS was under 300 bpm or if preparations were arrhythmic. Agonist doses were administered every 4 minutes using Gilson pipettes into the organ baths, with the tips submerged in the PSS. Atrial preparations were pre-treated with 1 µM Metoprolol 1 hour before the first dose of PE to avoid any interference from β-adrenoceptors. Inhibitors were administered 30 minutes prior to the first dose of PE, allowing sufficient time for these pharmacological agents to penetrate the multicellular preparations.

#### Neonatal myocytes isolation

Neonatal rat atrial myocytes (NRAMs) were isolated using a modified protocol from (Burton, Klimas et al. 2015, Burton, Tomek et al. 2020). Hearts were isolated from 3-day-old Sprague Dawley rat pups, culled by cervical dislocation followed by exsanguination. Following dissection, atria were separated from the ventricles, transferred to 2ml Dulbecco’s Modified Eagle’s Medium (DMEM) and cut into 1-2 mm^3^ pieces. Atrial myocytes were enzymatically isolated by a series of enzymatic digestions; in trypsin (1 mg/ml, Merck, UK, rocked at 4°C for 2 hours) followed by collagenase (type IV, 1mg/ml, Merck, UK). For the collagenase digestions, the trypsin was replaced by 4ml collagenase solution and tissue was gently triturated using a plastic pipette for 1 minute. The tissue plus collagenase solution was then stirred gently in a bath at 37 °C for 2 minutes and the first supernatant was discarded. A further 4ml collagenase solution was added to the tissue pellet and triturated for 1 minute using a wide bore pipette. This solution was then stirred gently in a bath at 37°C for 2 minutes and the supernatant (4ml) added to 3ml HBSS and stored in a 15ml centrifuge tube. This process was repeated a further 3 times to produce a total of 4×7ml cell suspensions. Tubes were centrifuged at 2000 rpm for 8 minutes. Supernatant was then removed, and the cells were resuspended in 2ml cardiomyocyte plating media (CPM: 85% DMEM, 17% M199, 10% horse serum, 5% FBS, 1% penicillin/streptomycin) before being centrifuged at 1000 rpm for 10 minutes. All samples were then strained using a cell strainer and pooled into a single 50ml centrifuge tube. Isolated cells were pre-plated and incubated at 37°C (95% O_2_, 5% CO_2_) for 1 hour to allow separation of fibroblasts. Supernatant containing suspended atrial myocytes was then removed and cell density measured using a haemocytometer and trypan blue. Myocytes were seeded onto 24mm laminin (40μg/ml, Merck, UK) coated glass coverslips in 35mm 6-well plates at a density of 15000 cells per ml in CPM. Cell media was changed every 48 hours. NRVMs were isolated as previously described (Zaccolo and Pozzan 2002).For isolation of neonatal mouse atrial myocytes (NMAMs) from WT or DKO C57BL/6, a similar protocol was followed. Enzymatic digestion was shorten du to smaller size and less amount of tissue. Enzymatic digestions: in trypsin rocked at 4 °C for 1.5 hours instead of 2 h. The collagenase digestions, atria were stirred gently in a bath at 37 °C for 1 minute instead of 2 minutes.

#### Adult mouse atrial isolation

WT or DKO C57BL/6 mice hearts were dissected and washed in heparin-containing PSS (20 IU per mL) on a Langendorff setup for retrograde perfusion via the aorta. The heart was fist perfused in a modified Tyrode (recipe in 1.10 Solutions, page) gassed with 95% O_2_/5% CO_2_ to maintain a pH of 7.4 at 37°C for 1 minute. Solution was switched to a digestion solution: the modified Tyrode above containing 100 µM CaCl_2_ and 0.02 mg/ml Liberase™ TH (Roche, Penzberg, Germany) and no EGTA. After 25 minutes of enzymatic digestion, the heart was removed. The atria were separated from ventricles in a dissection bath, the tissue was cut into small pieces (around 2×2mm) followed by 1 minute of gentle trituration and stored in a high potassium medium SANKB (in mM: KCl 70, MgCl_2_ 5, K^+^ glutamine 5, taurine 20, EGTA 0.04, succinic acid 5, KH_2_PO_4_ 20, HEPES 5, glucose 10; pH to 7.2 with KOH) at 4°C. Myocytes were left for 30 minutes before fixing for imaging or functional experiments (Ca^2+^ transient recordings), and were used up to 4 hours.

#### CaT mouse atrial

Single cell isolated atrial myocytes were incubated with 3 µM Fluo-5F-AM (Invitrogen™) in SANKB (recipe in 1.10 Solutions, page) for 10 minutes at 37°C in the dark. Cells were then plated to a glass coverslip for 10 minutes to adhere before imaging. Carbon fibre electrodes were used to field-stimulate Ca^2+^ transients at 1Hz. All recordings were at 37°C under gravity-fed superfusion of Tyrode (recipe in 1.10 Solutions, page). Cells were visualised using a Zeiss Axiovert 200 with attached Nipkow spinning disk confocal unit (CSU-10, Yokogawa Electric Corporation, Japan). Excitation light, transmitted through the CSU-10, was provided by a 488-nm diode laser (Vortran Laser Technology Inc., Sacramento, CA). Emitted light was passed through the CSU-10 and collected by an iXON897 EM-CCD camera (Oxford Instruments, UK) at 60 frames per second with 2×2 binning (pixel size = 0.66667 µm^2^). To avoid dye bleaching, video recordings were 10-20 seconds long (between 1000-2500 images) every minute to not continually be exposed to 488nm light. Ca^2+^ transients were measured using regions of interests (ROIs) in Andor iQ software (version 1.7) to record average whole cell fluorescence.

#### FRET

FRET imaging experiments were performed on day 4 of culture and 24 h after infection of the neonatal myocytes with adenovirus carrying EPAC-S^H187^ cytosolic biosensor (Klarenbeek, Goedhart et al. 2015) at a multiplicity of infection of 1000 virus particles per cell. During experiments, cells were maintained at room temperature in a modified Ringer solution (recipe in 1.10 Solutions, page). Dynamic measurements were done using an inverted microscope attached to a cool SNAP HQ2 camera and an optical beam splitter (Photometrics) for simultaneous recording of YFP and CFP emissions. FRET changes were measured as changes in the background-subtracted 480nm/545nm fluorescence emission intensity on excitation at 430 nm. For recording, cells were left to settle for 4 minutes before any additions of pharmacological agents. Saturation levels were ensured by addition of 10μM of forskolin that activates ACs and increases intracellular levels of cAMP and 100μM of 3-isobutyl-1-methylxanthine (IBMX) a competitive non-selective phosphodiesterase inhibitor which raises intracellular cAMP, activates PKA was added for maximum response. The number of technical and biological replicates is indicated in the figure legends.

#### hiPSC-aCM culture

Both hiPSC-aCM lines used for all experiments were derived from healthy donors. In particular, one from a female donor with no diagnosed diseases (https://hpscreg.eu/cell-line/TMOi001-A, purchased from Thermo Fisher Scientific) (Calamaio, Serzanti et al. 2023) and the other, previously characterized and published from a 62-year-old male not affected by AF or other cardiac pathologies (Benzoni, Campostrini et al. 2020). Approved protocols granted by the Ethical Committee of Brescia (protocol number 1737) and a written consent obtained from the patients, in agreement with the declaration of Helsinki. hiPSC-aCM lines were maintained on human Biolaminin 521 LN-coated dishes in TeSR-E8™ medium (Thermo Fisher Scientific). Cardiac differentiation was carried out by monolayer culture on Matrigel^®^ hESC-qualified Matrix (Corning, Corning, NY, USA) coated dishes using the PSC Cardiomyocytes Differentiation Kit (Thermo Fisher Scientific), following manufacturer instructions. To induce atrial differentiation, an adaptation of the method previously described (Devalla, Schwach et al. 2015) was used. Briefly, cells were treated with 1 μmol/L atRA (Sigma) starting from day 3 of differentiation, and the medium was changed every day until day 7. Then, cells were maintained in Cardiomyocytes Maintenance Medium (Thermo Fisher), changing the medium every other day. On the 21st day of differentiation, hiPSC-aCM culture was enriched using the PSC-Derived Cardiomyocyte Isolation Kit (Miltenyi Biotech) following manufacturer instructions, which allows magnetic separation with highly specific CM surface markers. Immediately after magnetic separation, iPSC-CMs were collected and cryopreserved in liquid nitrogen. For the experiments, atrial differentiated hiPSC-aCM were thawed on Matrigel-coated dishes and maintained at 37°C (95% O_2_, 5% CO_2_) with Cardiomyocyte Maintenance Medium with B-27 Supplement 50X and penicillin/streptomycin 100X (Thermo Fisher Scientific). Cells were culture until 28 days before running FRET or live imaging.

#### Drug information

Multiple pharmacological agents were used to inhibit the NAADP pathway at different steps and action cites. Bafilomycin A1 was administered at 100 nM, and acts as a vacuolar H+- ATPase inhibitor (Macgregor, Yamasaki et al. 2007, Collins, Bayliss et al. 2011). Trans-Ned-19, a TPC blocker and inhibits the Ca^2+^ signal was using at 1 µM (Pitt, Funnell et al. 2010). SG-094 (10 μM) is a specific TPC2 blocker that inhibits the Ca^2+^ release (Du, Guan et al. 2022) SAN4825 (10 μM) is an ADP-ribosyl cyclase inhibitor that reduces NAADP synthesis (Kannt, Sicka et al. 2012). BZ-194 is a small molecule that inhibits NAADP-mediated Ca^2+^ signalling administered at 500 μM. TPC2-A1-N (10 μM) is a Ca^2+^-permeable agonist of TPC2, and was administered to mimic the physiological actions of NAADP (Gerndt, Chen et al. 2020). DMSO (1:1000; vehicle) was added under control conditions to replicate addition of drugs that are diluted in DMSO.

##### Quantification and statistical analysis

For atrial preparations, [agonist] vs. response (three parameters) was used to form graphs and to calculate EC_50_s and maximum responses using Prism v10 software (GraphPad, CA, USA). Normalised data was used to compare responses. Maximum responses are presented as mean ± SD of recorded values, other than dose-response curve maximums and EC_50_ which are given as mean ± 95 % confidence interval (CI) of best-fit value. For FRET analysis, data was normalised and are presented as mean ± SD of recorded values.

Data was normalised and are presented as mean ± standard error of the mean (Macquaide, Tuan et al.) of recorded values. FRET Datasets in this manuscript were tested for normality distribution, using Shapiro Wilk normality test. If datasets passed normality test multiple-group comparison, a one-way analysis of variance (ANOVA) followed by multiple comparison testing was performed or for comparison between two groups, an unpaired, two-tailed Student’s t-test was used. If not, datsets were analysed using Kruskal Wallis followed by Dunn’s multiple comparison or non-parametric Mann-Whitney U test. Datasets are presented as mean ± SEM. Differences were considered statistically significant at values of P < 0.05, *P<0.05, ** P<0.01, *** P<0.001, **** P<0.0001. Statistical methods used are given in the legend of each figure. Statistical analyses were performed using Prism 10 (GraphPad, CA, USA). Differences were considered statistically significant at values of P < 0.05.

##### Data Accessibility

All relevant data can be shared by contacting the corresponding author.

## Supplemental Table, Videos and Figures

**Supplementary video 1 (relating to figure 2): WT HREM** Supplemental movie shows a 3D volume-rendered model of a wild type adult mouse heart generated from HREM data using Horos software.

**Supplementary video 2 (relating to figure 2): DKO HREM** Supplemental movie shows a 3D volume-rendered model of a double knockout AC1/AC8 type adult mouse heart generated from HREM data using Horos software.

## Notes

### Competing Interest Statement

The authors have declared no competing interest.

